# KRAB-zinc-finger proteins regulate endogenous retroviruses to sculpt germline transcriptomes and genome evolution

**DOI:** 10.1101/2023.06.24.546405

**Authors:** Kai Otsuka, Akihiko Sakashita, So Maezawa, Richard M. Schultz, Satoshi H. Namekawa

## Abstract

As transposable elements (TEs) coevolved with the host genome, the host genome exploited TEs as functional regulatory elements. What remains largely unknown are how the activity of TEs, namely, endogenous retroviruses (ERVs), are regulated and how TEs evolved in the germline. Here we show that KRAB domain-containing zinc-finger proteins (KZFPs), which are highly expressed in mitotically dividing spermatogonia, bind to suppressed ERVs that function following entry into meiosis as active enhancers. These features are observed for independently evolved KZFPs and ERVs in mice and humans, i.e., are evolutionarily conserved in mammals. Further, we show that meiotic sex chromosome inactivation (MSCI) antagonizes the coevolution of KZFPs and ERVs in mammals. Our study uncovers a mechanism by which KZFPs regulate ERVs to sculpt germline transcriptomes. We propose that epigenetic programming in the mammalian germline during the mitosis-to-meiosis transition facilitates coevolution of KZFPs and TEs on autosomes and is antagonized by MSCI.

## Introduction

Transposable elements (TEs) are mobile genetic elements that account for a large fraction (∼40-50%) of the mammalian genome (Koito and Ishizaka 2013; Trono 2015). Retrotransposons, a major group of TEs, work by a copy-and-paste mechanism using an RNA intermediate, causing DNA damage in the genome and leading to genome instability. In response to TEs, the host genome coevolved various defense mechanisms to silence and control retrotransposons (Zamudio and Bourc’his 2010; Elbarbary et al. 2016). Such mechanisms include DNA methylation, histone modifications, and piRNA pathways, all of which tightly control expression of TEs in the germline (Zamudio and Bourc’his 2010; Bao and Yan 2012; Di Giacomo et al. 2013; Crichton et al. 2014; Fu and Wang 2014; Ku and Lin 2014). However, TEs need to be expressed in the germline to propagate, and the host genome exploits TEs as functional regulatory elements to acquire reproductive fitness. Thus, as TEs coevolved with the host genome, the germline arose as the prominent battlefield to maximize mutual fitness (Zamudio and Bourc’his 2010; Elbarbary et al. 2016). Nevertheless, there remain major questions about how the activities of TEs are regulated, and TEs coevolved with the host in the germline.

Many TE-derived sequences function as gene regulatory elements, such as promoters and enhancers, to drive tissue- and cell-type-specific gene expression (Peaston et al. 2004; Pi et al. 2004; Rebollo et al. 2012; Erwin et al. 2014; Friedli and Trono 2015; Garcia-Perez et al. 2016; Thompson et al. 2016; Chuong et al. 2017; Huang et al. 2017). Genome-wide studies demonstrate that a significant portion of TF-binding sites is derived from TEs (Rebollo et al. 2012; Sundaram et al. 2014). Comprising ∼10% of the mammalian genome, endogenous retroviruses (ERVs) are subfamilies of TEs and remnants of retroviruses integrated into the genome (Waterston et al. 2002). Most ERVs in the mammalian genome are truncated but remain as long terminal elements (LTR). Testis-specific expression of ERVs in humans and mice was initially reported over 40 years ago (Del Villano and Lerner 1976), and recent studies reveal regulatory functions for TEs in male meiosis. These functions encompass post-transcriptional regulation of mRNA and long noncoding RNAs (lncRNAs) via the piRNA pathway (Watanabe et al. 2015), as well as promoter functions for ERVs that drive lncRNA expression (Davis et al. 2017). Our recent study demonstrated that ERVs function as active enhancers to drive expression of species-specific germline genes during the mitosis-to-meiosis transition (Sakashita et al. 2020), a critical developmental transition during which both the chromatin and epigenome are dramatically changed (Sin et al. 2015; Maezawa et al. 2018; Alavattam et al. 2019; Patel et al. 2019; Wang et al. 2019). Largely unknown, however, is how the activities of ERVs are dynamically regulated during spermatogenesis.

To determine the regulatory mechanism of ERVs enhancers to generate germline transcriptomes, here we focus on rapidly evolved ERVs and Krüppel-associated box (KRAB) domain-containing zinc-finger proteins (KZFPs), a group of transcription factors (TFs) that underwent coevolution to bind and suppress expression of ERVs (Tadepally et al. 2008; Ecco et al. 2017; Yang et al. 2017; Bruno et al. 2019; Wolf et al. 2020; Senft and Macfarlan 2021). KZFPs bind DNA using tandem arrays of C2H2 zinc finger domains, each of them recognizes two to four specific nucleotides, thereby recognizing specific DNA sequences with high affinity (Patel et al. 2018). There are hundreds of ERVs and KZFPs in mammals, with many pairwise functional interactions between them (Imbeault et al. 2017), suggesting an evolutionary arms race between the host genome and viral-derived elements (Ecco et al. 2017; Yang et al. 2017; Bruno et al. 2019; Senft and Macfarlan 2021). Although previous studies revealed that KZFPs function to suppress ERV activity in cultured cells (Imbeault et al. 2017; Kumar et al. 2020), *in vivo* functions of KZFPs remain elusive (Wolf et al. 2020; Pontis et al. 2022). Here, we report a mechanism by which KZFPs regulate the activity of ERV-derived enhancers to sculpt germline transcriptomes necessary for spermatogenesis in mice and humans. We show how KZFPs and TEs coevolved in the epigenetic programming at the mitosis-to-meiosis transition and how they were impacted by meiotic sex chromosome inactivation (MSCI), an essential process in the male germline (Turner 2015; Alavattam et al. 2021). Our study reveals meiosis as the battlefield of an arms race between the host genome and retroviruses in their coevolution.

## Results

### Expression of KZFPs and ERVs dynamically changes during spermatogenesis in mice

Because KZFP family proteins rapidly evolved in mammals and contributed to the evolution of gene regulatory networks in mammals (Imbeault et al. 2017; Senft and Macfarlan 2021), we first evaluated the sequencing similarity of coding KZFP genes of mouse and human KZFPs among various species of mammals. Notably, a subset of mouse KZFPs was only present in mice, and the other subset of mouse KZFPs were unique to rodents, including rats and Chinese hamsters, and others shared low sequence homologies to their counterparts in other mammals (Fig. 1A). In addition to these rapidly evolved KZFPs, a small group of KZFPs was highly conserved among mammals (Fig. 1A). The same tendency was also observed for human KZFPs, including human-specific, primate-specific, lowly conserved, and highly conserved KZFPs (Fig. 1A). These analyses, together with a recent review (Senft and Macfarlan 2021), clarify that the rapid evolution of KZFPs and a subset of KZFPs are unique to evolutionary orders, such as rodents and primates.

**Figure 1.**
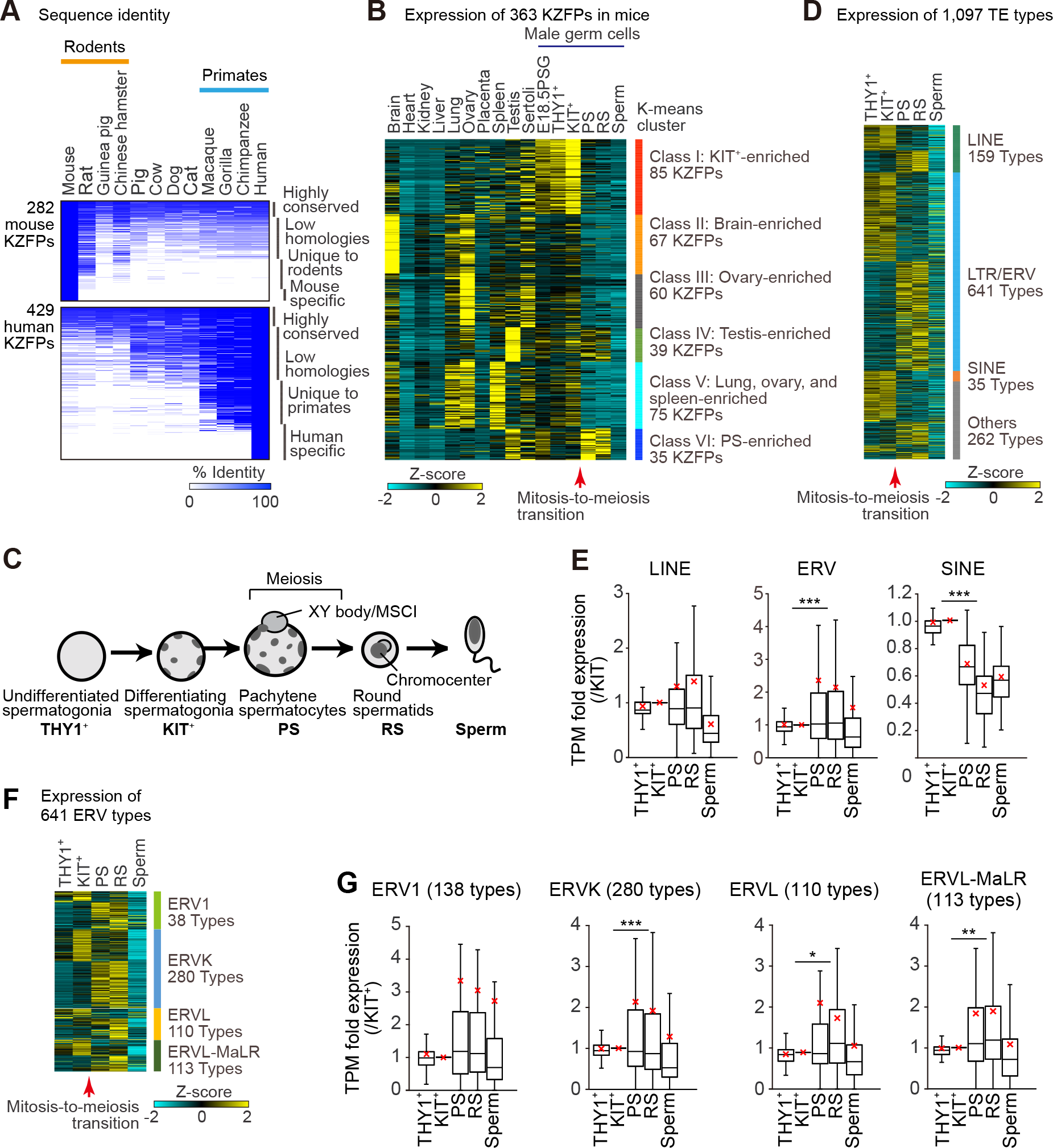
Expression of KZFPs and TEs during mouse spermatogenesis. **A**, Sequence identities of 282 murine KZFPs (top) and 429 human KZFPs (bottom) among 12 mammalian species. **B**, A heatmap showing a k-means clustering analysis of expression (RNA-seq data) for all protein-coding KZFPs in various tissues and testicular cell types in mice. E18.5PGC: prospermatogonia from E18.5 testis, THY1^+^: THY1^+^ undifferentiated spermatogonia, KIT^+^: KIT^+^ differentiating spermatogonia, PS: pachytene spermatocytes, RS: round spermatids. **C**, Schematic of mouse spermatogenesis and the five representative stages. **D**, A heatmap showing the expression (RNA-seq data) of all TE types in male germ cells at the representative five stages. **E**, Relative expression (RNA-seq data) of each TE class during spermatogenesis. The values of KIT^+^ were set to 1. The red crosses represent the average values. ANOVA followed by Tukey-HSD test. ***: p<0.001. **F**, A heatmap showing the expression (RNA-seq data) of all ERV types in mouse spermatogenesis. **G**, Relative expression (RNA-seq data) of each ERV subclasses during spermatogenesis. The values of KIT^+^ were set to 1. ANOVA followed by Tukey-HSD test. *: p<0.05, **: p<0.01, ***: p<0.001.

Because rapid gene evolution is a feature of germline genes expressed in testes (Soumillon et al. 2013), we suspected that KZFPs are dynamically expressed in the male germline in testes compared to other tissues. Accordingly, we analyzed expression of 363 mouse protein-coding KZFPs using RNA sequencing (RNA-seq) data sets from 9 tissues, testicular somatic Sertoli cells, and represented cell types in the male germline (Fig. 1B). For the analysis of male germ cells, we examined THY1^+^ undifferentiated spermatogonia, which contain a stem cell population; THY1^+^ differentiated spermatogonia, which committed spermatogenic differentiation; pachytene spermatocytes (PS), which is amid meiotic prophase I; and haploid round spermatids (RS: Fig. 1C). We were able to categorize the 363 murine protein-coding KZFPs into six major classes based on non-hierarchical k-means clustering analysis (Fig. 1B, Supplemental Table 1). Class I KZFPs include 85 KZFPs, which were highly expressed in mitotically dividing male germ cells but suppressed after the mitosis-to-meiosis transition. We also detected classes of KZFPs that were highly enriched in specific tissues (Class II: brain-enriched 67 KZFPs; Class III: ovary-enriched 60 KZFPs, which were also highly expressed in Sertoli cells, Class IV: testis-enriched 39 KZFPs, and Class V: lung, ovary, and spleen-enriched 75 KZFPs). Classes II, III, and V were downregulated at the mitosis-to-meiosis transition. In contrast, Class VI KZFPs, encompassing 35 PS-enriched KZFPs, were upregulated when germ cells reached the pachytene stage. Thus, nearly 90% of KZFPs undergo dynamic expression changes at the mitosis-to-meiosis transition in the male germline.

We next examined changes in KZFP expression during different stages of meiosis using previous single-cell RNA-seq data (Hermann et al. 2018b) (Fig. S1A). Class I, II, III, and V KZFPs were downregulated by the leptotene and zygotene stages of meiotic prophase I, whereas expression of Class VI KZFPs was observed from the pachytene stage, i.e., the transition occurs during early meiotic prophase I. Although the expression patterns of these KZFPs are dynamic, GO term analysis revealed that all classes of KZFPs are enriched with gene functions that share common features, such as negative regulation of transcription and transcriptional repressor activity (Fig. S1B). This result suggests that transcriptional repression is a major function of each class of KZFPs.

Because KZFPs are thought to regulate expression of TEs (Imbeault et al. 2017; Senft and Macfarlan 2021), we next evaluated the dynamics of TE expression at representative stages of spermatogenesis by analyzing the expression of each unique TE copy. We detected 486,155 unique expressed TE copies in spermatogenesis (Fig. S1C), and these unique copies were categorized into 1,097 TE types, including 159 types of long interspersed nuclear elements (LINEs), 641 types of LTR/ERVs (abbreviated as ERVs hereafter), and 35 types of short interspersed nuclear elements (SINEs: Fig. 1D). Consistent with our recent report (Sakashita et al. 2020), expression of TEs dynamically changed at the mitosis-to-meiosis transition (Fig. 1D). Of note, overall expression of ERVs was significantly upregulated in PS compared to KIT^+^ spermatogonia (Fig. 1E), whereas expression of SINEs was largely downregulated at the mitosis-to-meiosis transition (Fig. 1D, E). The overall upregulation of ERVs in PS mirrors the overall downregulation of KZFPs in PS, suggesting a mechanistic linkage. Thus, we further investigated the expression of four representative ERV families, ERV1, ERVK, ERVL, and ERVL-MaLR. Among them, expression of ERVK, ERVL, and ERVL-MaLR was significantly increased at the mitosis-to-meiosis transition, whereas ERV1 expression did not change (Fig. 1F, G). These results suggest that ERVs, especially expression of ERVKs, ERVLs, and ERVL-MaLRs, are generally suppressed at the pre-meiotic stage and are then expressed in the meiotic stage.

### KIT^+^ spermatogonia-enriched KZFPs target specific TEs in mice

Because KZFPs are highly expressed in pre-meiotic cells and TEs, particularly ERVs, become active in meiotic cells, we hypothesized that premeiotic KZFPs suppress expression of TEs and downregulation of KZFPs leads to activation of TEs in meiotic cells. To test this hypothesis, we first determined possible interactions between KZFPs and TEs by reanalyzing ChIP-seq data for mouse 61 KZFPs in cultured cell lines, including embryonic stem cells and embryonic carcinoma cells (Wolf et al. 2020). Because of the sequence specificity of KZFP binding—KZFP binding sites are considered conserved among various cell types (Imbeault et al. 2017; Wolf et al. 2020)—this strategy should identify common binding sites of KZFPs present in KIT^+^ spermatogonia. Among 61 KZFPs, 13 belong to KIT^+^ spermatogonia-enriched Class I KZFPs and consequently we examined their binding sites and determined their binding enrichment to TEs across the mouse genome. We first detected peaks for 13 KZFPs from the ChIP-seq data by following the criteria described previously (Wolf et al. 2020) (Fig. S2A). Among 13 KZFPs, several KZFPs, including GM14406, GM14401, and ZFP987, showed unique binding sites that are not shared by other members of 13 KZFPs (Fig. 2A). Next, we examined whether these peaks are enriched on TEs in the genome. Among 13 KZFPs, GM14406, ZFP600, REX2, GM14406, and ZFP990 showed significant enrichment at all TE loci compared to random loci (Fig. S2B). Enrichment to TEs, however, is class-specific; GM14406 on LINEs (Fig. 2B), ZFP989 and ZFP987 on ERVs (Fig. 2C), and GM14393, ZFP990, and ZFP986 on SINEs (Fig. 2D). These results demonstrate the target specificity of KIT^+^- enriched KZFPs.

**Figure 2.**
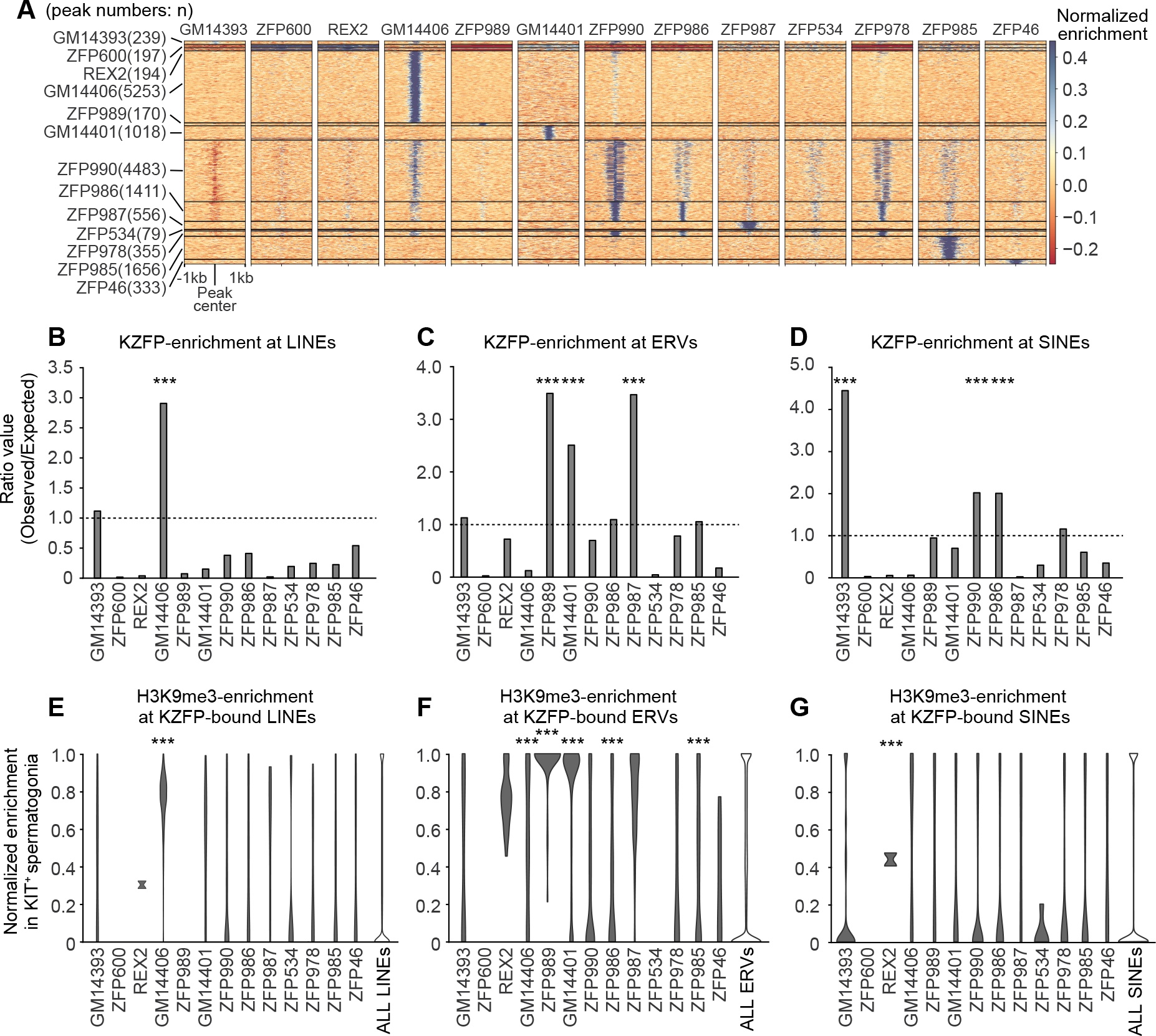
Binding preference of KIT^+^ spermatogonia-enriched KZFPs and the H3K9me3 enrichment. **A**, Heatmap of the binding profiles of KIT^+^ spermatogonia-enriched KZFPs. The numbers in parentheses represent the numbers of detected peaks. **B, C, D**, Binding preference of KZFPs at LINEs (**B**), ERVs (**C**), and SINEs (**D**). The bar plot represents the ratio value of the observed numbers of peaks versus the theoretically expected numbers of peaks. Binomial Test. ***: p<0.001. **E**, **F**, **G,** Normalized H3K9me3 enrichment at each KZFP-bound LINEs (**E**), ERVs (**F**), and SINEs (**G**) compared to enrichment at all interspersed copies in KIT^+^ spermatogonia. Mann-Whitney U-test. ***: p<0.001.

A general function of KZFPs is to repress transcription by inducing H3K9me3 to the binding loci (Schultz et al. 2002; Groner et al. 2010; Bruno et al. 2019). Therefore, we next examined if the KZFPs binding sites are enriched with H3K9me3 in KIT^+^ spermatogonia. In accordance with the KZFP enrichment on specific classes of TEs, we observed significant enrichment of H3K9me3 at GM14406- bound LINEs (Fig. 2E) and at ZFP989-bound LTR/ERVs (Fig. 2F). Enrichment of H3K9me3 was also observed at other KZFP-bound LTR/ERVs, including GM14406, GM14401, ZFP986, and ZFP985 (Fig. 2F). Such H3K9me3 enrichment was not observed on KZFP-bound SINEs except REX2-bound SINEs (Fig. 2G), although REX2 does not preferentially associate with SINEs in the genome (Fig. 2D). These results suggest that KIT^+^-enriched KZFPs independently recognize specific classes of TEs and establish H3K9me3 at the target loci.

### Correlation between expression of target ERVs and adjacent genes in mice

To further elucidate how KIT^+^-enriched KZFPs regulate target ERVs, we focused on further analysis of 3 ERV-targeting KZFPs, ZFP989, Gm14401, and ZFP987, which showed significant enrichment at ERV loci (Fig. 2C, 3A). Because these KIT^+^-enriched KZFPs bind to specific ERVs that are enriched with H3K9me3 in KIT^+^ spermatogonia, we suspected that the target ERVs would be activated as gene regulatory elements when expression of these KZFPs is repressed in the meiotic stage. To test this possibility, we first compared the H3K9me3 level at the target ERVs at the mitosis-to-meiosis transition. At the KIT^+^-enriched KZFP (ZFP989, Gm14401, and ZFP987) target 1,170 ERV loci (Fig. 3A), H3K9me3 was decreased from KIT^+^ spermatogonia and PS (Fig. 3B). The reduction of H3K9me3 was confirmed in track views of representative ERV loci that are targets of KIT^+^-enriched KZFPs (Fig. S3A). Of note is that H3K27 acetylation (H3K27ac), a marker of active enhancers, was increased at these ERV loci from KIT^+^ spermatogonia and PS (Fig. 3C). Further examination of substages during meiotic prophase I confirmed the progressive reduction of H3K9me3 and gain of H3K27ac at these ERV loci (Fig. S3B, C). These results suggest that H3K9me3 KIT^+^-enriched KZFP-target ERVs are released from KZFP-mediated silencing and gain a feature of active enhancers at the mitosis-to-meiosis transition.

**Figure 3.**
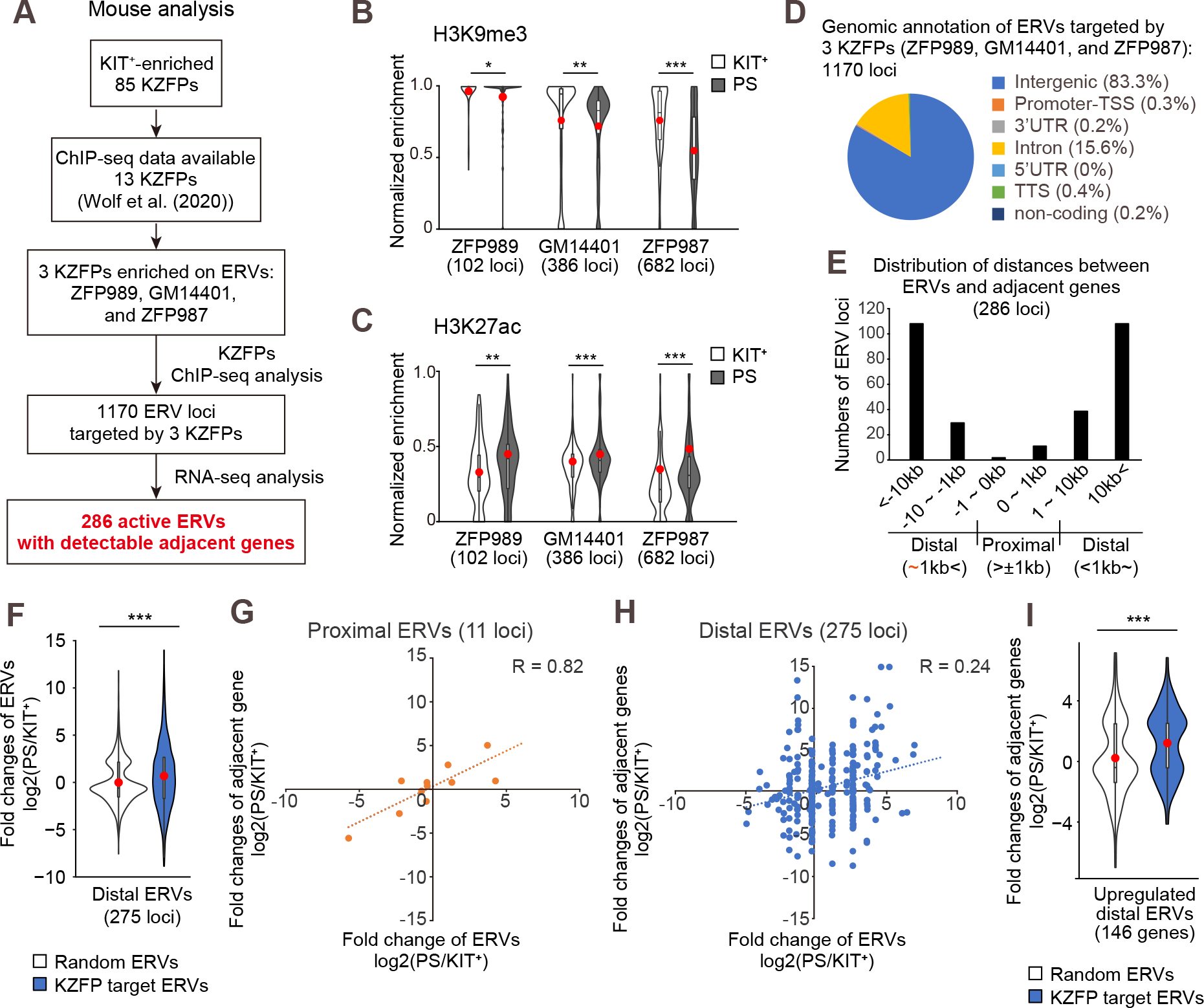
KZFPs target and suppress ERVs until the mitosis-to-meiosis transition. **A**, Flowchart of analyses to identify target ERVs. **B,C,** Normalized enrichment of H3K9me3 (**B**) and H3K27ac (**C**) at the target ERV loci in KIT^+^ and PS. Mann-Whitney’ s U-test. ***: p<0.001, **: p<0.01. *: p<0.05. **D**, Genome annotation of the ERV loci targeted by ZFP989, GM14401, and ZFP987. Each proportion is shown in parentheses. **E**, Distances between ERVs targeted by ZFP989, GM14401, and ZFP987 and their adjacent genes (Proximal: <1kb, Distal: 1kb<). **F**, Fold expression change of target distal ERVs at the mitosis-to-meiosis transition (PS/KIT^+^), Student’ s t-test. ***: p<0.001. **G**, **H**, Correlation between the expression of the adjacent gene and proximal ERVs (**G**), or distal ERVs (**H**) at the mitosis-to-meiosis transition (PS/KIT^+^). **I**, Fold expression change of the genes adjacent to upregulated distal ERVs at the mitosis-to-meiosis transition (PS/KIT^+^). Student’ s t-test. ***: p<0.001.

Consistent with these epigenomic features, the KZFP-target 1,170 ERVs are mostly located in intergenic or intronic regions (Fig. 3D). Given our previous finding that a subset of ERVs is activated as enhancers in meiosis to drive spermatogenic gene expression (Sakashita et al. 2020), we hypothesized that these ERVs functions as enhancers. To test this hypothesis, from the 1,170 ERVs potentially regulated by KZFPs, we further selected 286 ERVs, for which their transcripts and the adjacent gene transcripts were detected either in KIT^+^ spermatogonia and PS by RNA-seq (Fig. 3A). Among these 286 active ERV loci, 11 loci were located within 1 kb of adjacent genes (termed proximal ERVs), whereas 275 ERVs loci were located more than 1 kb from adjacent genes (termed distal ERVs: Fig. 3E). These 275 distal ERVs were significantly upregulated from KIT^+^ to PS compared to randomly selected ERVs (Fig. 3F). Both proximal and distal ERVs showed positive correlations between upregulation of ERVs and upregulation of their adjacent genes from KIT^+^ to PS (Fig. 3G, H). Accordingly, adjacent genes of upregulated distal ERVs were upregulated in PS (Fig. 3I), suggesting that distal ERVs act as enhancers for adjacent genes.

Together, we concluded that KIT^+^-enriched KZFPs suppress the activity of ERVs and that the downregulation of KZFPs activates ERV enhancers in meiotic cells in mice.

### Knockdown of KIT^+^-enriched KZFPs results in derepression of target ERVs and adjacent genes

Next, we sought to validate the functions of KIT^+^-enriched KZFPs in suppressing meiotic ERVs. To this end, we performed siRNA KZFP knockdown experiments for KIT^+^-enriched KZFPs using embryonic stem cells (ESCs) in which meiotic ERVs enhancers and germline genes are not active and determined whether meiotic ERVs, which are targets of these KIT^+^-enriched KZFPs, and their adjacent genes were derepressed. ZFP987 binds and suppresses a type of meiotic ERV, *ERVB4_1C-LTR*, and their adjacent germline genes, *Orf19* and *Vmn2r100* (Fig. 4A). Targeting *Zfp987* reduced expression of *Zfp987* and *ERVB4_1C-LTR, Orf19,* and *Vmn2r100* were derepressed (Fig. 4B). In line with this observation, an overall reduction of H3K9me3 was observed at both *Orf19* and *Vmn2r100* genes loci at the KIT^+^ to PS transition (Fig. 4C, D). This result suggests that ZFP987 suppresses *ERVB4_1C-LTR* via H3K9me3 in KIT^+^, and *ERVB4_1C-LTR* and target genes are activated in PS after the loss of ZFP987. Thus, the result observed in siRNA knockdown experiments likely reflect *in vivo* events in the mitosis-to-meiosis transition. We also performed siRNA knockdown experiments for another KIT^+^-enriched KZFP, *Zfp989,* which targets a type of ERV, *IAPLTR1a*, adjacent to the *Ube3a* gene (Fig. 4E). siRNA knockdown of *Zfp989* resulted in derepression of *IAPLTR1a* and the *Ube3a* gene (Fig. 4F). In the KIT^+^ to PS transition, the reduction of H3K9me3 was observed at ERV loci adjacent to the *Ube3a* gene locus (Fig. 4G).

**Figure 4.**
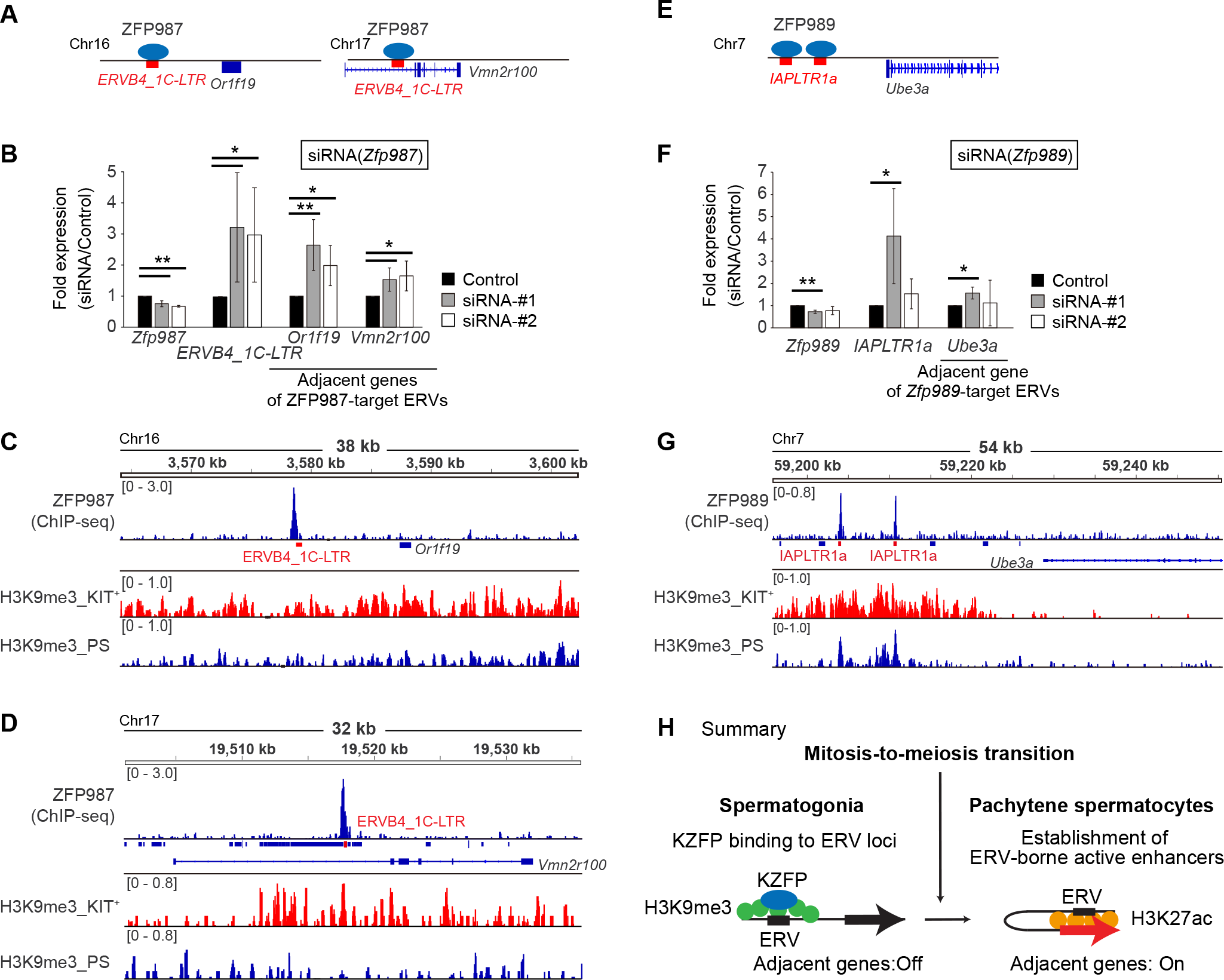
Knockdown of KIT^+^-enriched KZFPs results in derepression of ERVs and their adjacent genes. **A, E** Schematic of ERV enhancers and their target loci. **B, F**. Result of qRT-PCR for KZFP, targeted ERV, and ERV-adjacent genes under *Zfp987* knockdown (**b**) and *Zfp989* knockdown (**f**). Data values for controls were normalized to 1.0. Student’ s t-test. **: p<0.01, *: p<0.05. n=4. **C, D, G**. Track views of KZFP and H3K9me3 enrichment in KIT^+^ and PS at the target ERV loci. Genomic regions around target genes (*Or1f19* (**c**), *Vmn2r100* (**d**), and *Ube3a* (**g**)) are shown. Target ERVs are represented as red-colored letters and quadrangles. **H**. Summary. In spermatogonia, spermatogonia-enriched KZFPs bind and suppress ERVs that later function as enhancers. At the ERV loci targeted by spermatogonia-enriched KZFPs, after entering meiosis, KZFPs were displaced, and H3K9me3 decreased, while H3K27ac increased to become active enhancers.

Together, we conclude that KIT^+^-enriched KZFPs suppress target ERVs via H3K9me3, and the release from KZFP-mediated repression enables activation of ERV enhancers and their target genes, which takes place in the mitosis-to-meiosis transition in spermatogenesis (Fig. 4H).

### Human KZFPs contribute to ERV suppression in male germ cells at the mitotic stage

We next sought to examine whether human KZFPs, which underwent distinct evolution from mouse KZFPs, regulate expression of meiotic ERVs. Like mouse ERVs, human ERVs underwent rapid evolution in mammals. Some human ERVs act as enhancers to drive species-specific genes in meiosis (Sakashita et al. 2020), raising the possibility that KZFPs suppress meiotic ERVs in premeiotic cells in humans. To test this possibility, we reanalyzed previous RNA-seq data of human spermatogenic cells (Zhu et al. 2016).

Human KZFPs showed a drastic change in expression at the mitosis-to-meiosis transition from spermatogonia (SG) to primary spermatocytes (PriSC), which include pachytene spermatocytes (Fig. 5A). Human KZFPs further undergo a further change from PriSC to round spermatids (RS) (Fig. 5A). 469 human KZFPs were categorized into three classes, including 118 SG-enriched KZFPs (Class I), 211 PriSC-enriched KZFPs, and 141 RS-enriched KZFPs (Fig. 4A, Supplemental Table 2). Likewise, human TEs, including LINEs and ERVs undergo dynamic expression changes during spermatogenesis (Fig. 5B), consistent with our previous analysis (Sakashita et al. 2020). LINEs and ERVs were significantly upregulated from SG to PriSC (Fig. S4A). Of note, similar to mouse, human SINEs were predominantly expressed in SG but were suppressed in PriSC and RS (Fig. 5B, S4A).

**Figure 5.**
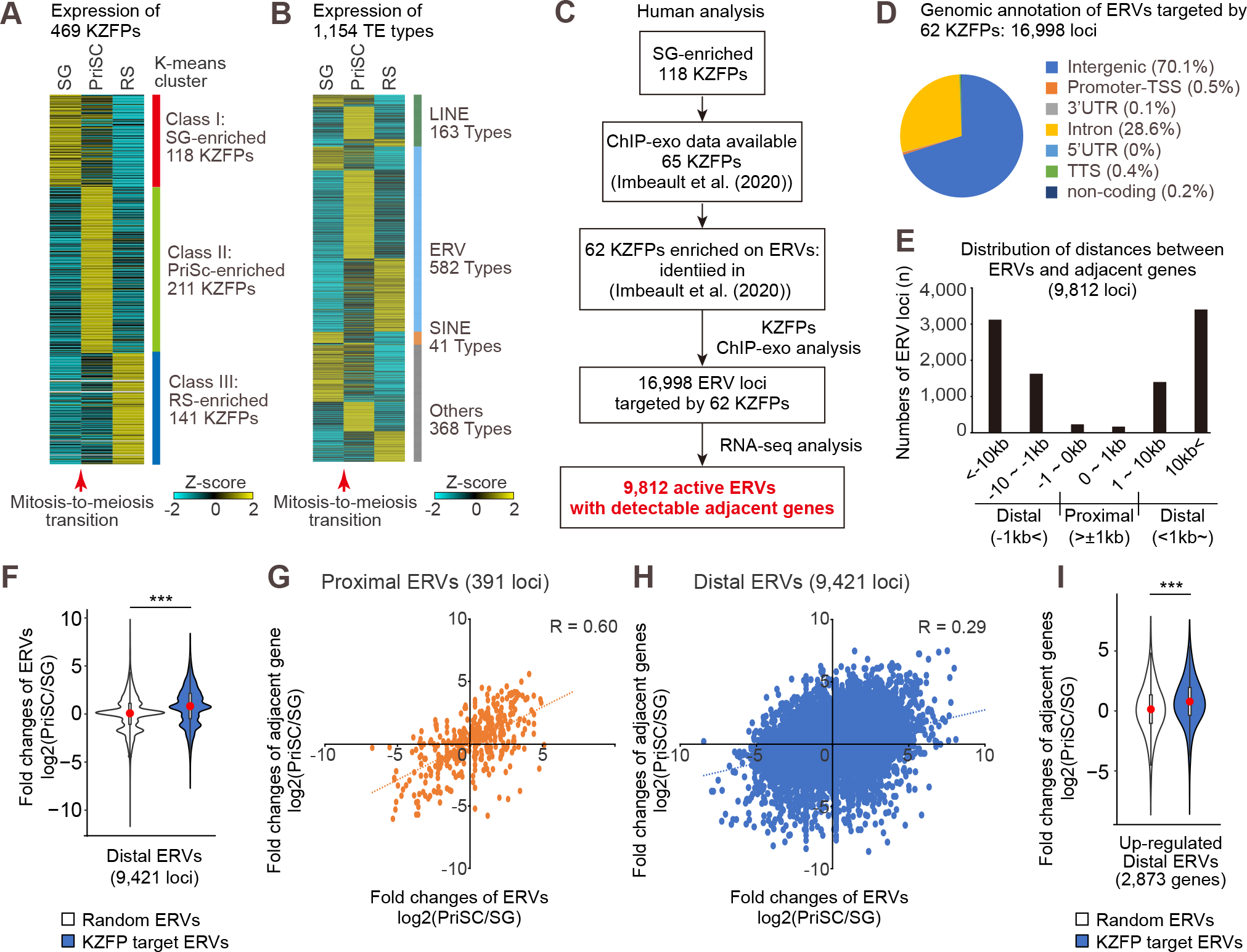
Human pre-meiotic KZFPs can act as suppressors of meiotic enhancer ERVs. **A**, Heatmaps showing a k-means clustering analysis of expression (RNA-seq data) for all protein-coding KZFPs in human testicular germ cells. **B**, Heatmaps showing the expression (RNA-seq data) of all TE types in human testicular germ cells. **C**, Flowchart of analyses to identify target ERVs. **D**, Genome annotation of the human ERV loci targeted by 62 human SG-enriched KZFPs. **E**, Distances between ERVs targeted by 62 human SG-enriched KZFPs and their adjacent genes (Proximal: <1kb, Distal: 1kb<). **F**, Fold-expression change of target distal ERVs at the mitosis-to-meiosis transition (PriSC/SG), Student’ s t-test. ***: p<0.001. **G, H**, Correlation between expression of the adjacent gene and proximal ERVs (G), or distal ERVs (H) at the mitosis-to-meiosis transition (PriSC/SG). **I**, Fold-expression change of the genes adjacent to upregulated distal ERVs at the mitosis-to-meiosis transition (PriSC/SG). Student’ s t-test. ***: p<0.001.

To examine the interactions between human KZFPs and TEs, we reanalyzed a previous data set of ChIP-exo (ChIP with lambda exonuclease and sequencing) experiments of 222 human KZFPs in HEK293T cells (Imbeault et al. 2017). Among 118 SG-enriched KZFPs, 65 KZFPs have ChIP-exo data, and 62 KZFPs are highly enriched on ERVs based on the previous analysis (Imbeault et al. 2017) (Fig. 5C). Using these data, we detected binding peaks that have varied frequencies of unique peaks that do not overlap with peaks of other KZFPs (Fig. S4B, C). These 62 KZFPs bind 16,998 ERV loci, 70.1 % of which are located in intergenic regions and 28.5% of which are located in introns (Fig. 5D).

Among 16,998 ERV loci targeted by SG-enriched KZFP, we further selected 9,812 ERVs, for which their transcripts and the adjacent gene transcripts were detected either in SG and PriSC in RNA-seq (Fig. 5C). Consistent with our prediction of their enhancer functions, the distances between human ERVs and their adjacent genes were mostly located more than 1 kb (9,421 loci) and therefore these ERVs were defined as distal ERVs (Fig. 5E). The distal ERVs showed increased expression from SG to PriSC (Fig. 5F). On the other hand, a small number of ERVs showed a distance < 1 kb from their adjacent genes (391 proximal loci: Fig. 5E), and their overall expression changes from SG to PriSC were comparable when compared to randomly selected loci (Fig. S4D). Both proximal and distal ERVs showed positive correlations between increased expressions of ERVs and their adjacent genes from SG to PriSC (Fig. 5G, H). Furthermore, adjacent genes of these upregulated ERVs from SG to PriSC were likewise upregulated (Fig. 5I, S4E). These results suggest that in SG, SG-enriched KZFPs bind and suppress expression of ERVs that later function as gene regulatory elements, such as promoters (proximal ERVs) and enhancers (distal ERVs).

### The mitosis-to-meiosis transition facilitates the coevolution of KZFPs and ERVs in mammals

Our analysis provides a molecular mechanism by which a group of KZFPs controls ERVs to regulate meiotic gene expression in mammals. Because KZFPs and ERVs rapidly evolved by the arms race between them (Imbeault et al. 2017; Senft and Macfarlan 2021), we determined the evolutionary history of spermatogonia-enriched KZFPs and their target ERVs that are activated in meiosis. In mice, 66 KIT^+^- enriched KZFPs rapidly evolved, and a similar trend was found in all KZFPs (Fig. 6A). However, when we compared the sequence identities between mouse and rat, 66 KIT^+^-enriched KZFPs were relatively well conserved compared to other KZFPs (Fig. 6B), a feature not observed between mouse and human (Fig. 6B), suggesting that the KIT^+^-enriched KZFPs could function in close species such as mouse and rat. Among mouse KIT^+^-enriched KZFPs, we especially focused on three ERV-targeting KZFPs, ZFP989, GM14401, and ZFP987, because they preferentially bound ERVs (Fig. 2C). These three KZFPs were identified as *Mus* genus-specific (Imbeault et al. 2017) and recognized a specific subset of *Mus* genus-specific ERVKs, such pairs include ZFP989-*IAPLTR1a_Mm*, GM14401-*IAPEz-int*, and ZFP987-

**Figure 6.**
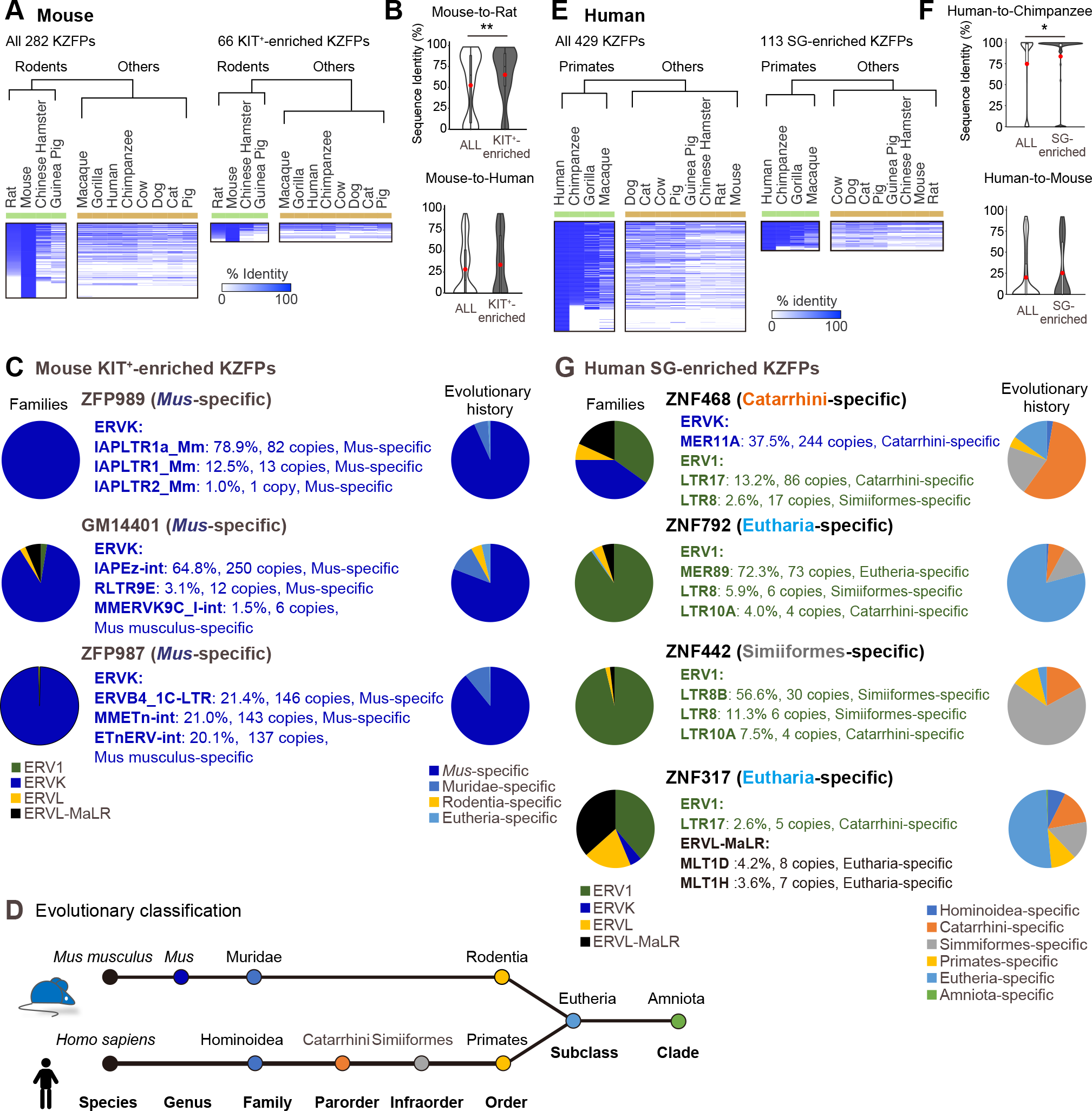
The evolutional aspects of KZFPs enriched in the mitotic spermatogonia. **A**, The heatmap represents the identity of all 282 murine KZFPs (left) and 66 KIT^+^-enriched KZFPs (right) compared among mammalian species. The dendrogram represents the result of the hierarchical analysis, which categorizes all species into two groups. **B**, The violin plot represents the comparison of the sequence identity between ALL KZFPs (white) and KIT^+^-enriched KZFPs (grey) between mice and rats (top) and mice and humans (bottom). Mann-Whitney’ s U-test. **: p<0.01. **C**, The left pie chart represents the proportion of subfamilies of target ERVs. The name of the ERV types shown in the middle represents the top 3 ERV types targeted by each KZFP. The right pie chart represents the proportion of evolutional ages of target ERVs. Evolutional history of each KZFP is shown in parentheses. **D**, Schematics of evolutional clades for mice (top) and humans (bottom). **E**, The heatmap represents the identity of all 429 human KZFPs (left) and 113 SG-enriched KZFPs (right) compared among mammalian species. The dendrogram represents the result of the hierarchical analysis, which categorizes all species into two groups. **F**, The violin plot represents the comparison of the sequence identity between ALL KZFPs (white) and SG-enriched KZFPs (grey) between humans and chimpanzees (top), and humans and mice (bottom). Mann-Whitney’ s U-test. *: p<0.05. **G**, Evolutional age of 4 representative human SG-enriched KZFPs and their target ERVs. The left pie chart represents the proportion of subfamilies of target ERVs. The name of the ERV types shown in the middle represents the top 3 ERV types targeted by each KZFP. The right pie chart represents the proportion of evolutional ages of target ERVs.

*ERVB4_1C-LTR* (Fig. 6C, D). The ERVK family showed the most significant upregulation after the mitosis-to-meiosis transition in mice (Fig. 1F), and indeed target ERVs of these three KZFPs are upregulated in PS (Fig. S5A). Thus, mouse spermatogonia-enriched KZFPs target newly-evolved ERVKs, which become active in meiosis.

We next examined evolutionary aspects of human KZFPs. Overall, 113 SG-enriched KZFPs have relatively high sequence homologies among primates and are less enriched with human-specific KZFPs (Fig. 5E). Consistent with this feature, SG-enriched KZFPs are relatively well-conserved between human and chimpanzee, whereas sequence identities are not so high between human and mouse (Fig. 6F). This feature is consistent with the evolutional feature of mouse KZFPs, and relatively similar spermatogonia-enriched KZFPs in closely related species may regulate expression of germline genes. We then focused on four representative SG-enriched KZFPs, which showed the most drastic decrease at the mitosis-to-meiosis transition (Fig. S4E). In contrast with the mouse KIT^+^-enriched KZFPs, which mainly targeted the ERVK family, human SG-enriched KZFPs targeted various ERV families with a bias towards the ERV1 and ERVL-MaLR families (Fig. 6G). The evolution of these four KZFPs corresponds to the evolution of target ERVs; Catarrhini-specific ZNF468 targeted Catarrhini-specific MER11A and LTR17, and Eutheria-specific ZNF792 and ZNF317 primarily targeted Eutheria-specific ERVs. Furthermore, Simiiformes-specific ZNF442 targeted Simiiformes-specific ERVs. These target ERVs were upregulated in PriSC (Fig. S5B). These results suggest the concomitant evolution of spermatogonia-enriched KZFPs and their target meiotic ERVs in mammals, which is consistent with the arms race between these KZFPs and ERV pairs being necessitated by the need to undergo the mitosis-to-meiosis transition in mammals.

### Meiotic KZFPs are not associated with ERV suppression in meiosis

In contrast to spermatogonia-enriched KZFPs, groups of KZFPs were highly upregulated both in mice and humans after the mitosis-to-meiosis transition. In mice, 35 Class VI KZFPs were highly expressed in PS (Fig. 1B), raising the possibility that they function in meiotic cells. Of note, these 35 KZFPs are relatively well conserved among mammals (Fig. S6A), and their human orthologs tend to be expressed in meiosis or in postmeiotic round spermatids (Fig. S6B). To characterize these PS-enriched KZFPs, we first analyzed the binding preference of five PS-enriched KZFPs whose ChIP-seq data were available (Fig. S6E). Among these five KZFPs, ZFP992, ZKSCAN17, and ZFP94 showed a significant enrichment at TE loci. ZFP992 and ZFP94 were significantly enriched at ERVs, and ZFP992, ZFP94, and ZFP457 were significantly enriched at SINEs, whereas none of the 5 KZFPs were preferentially bound to LINEs (Fig. S6F). However, these target ERVs and SINEs did not show expression changes at the mitosis-to-meiosis transition (KIT^+^ to PS: Fig. S6G, H), suggesting that PS-enriched KZFPs are not associated with suppression of target ERVs in meiosis in mice.

We next examined if human KZFPs highly expressed in meiosis are associated with suppression of target ERVs in meiosis. In contrast to mouse meiotic KZFPs, which are relatively well conserved in mammals (Fig. S6A), human PriSC-enriched KZFPs are not well conserved among mammals (Fig. S7A). Among 211 PriSC-enriched KZFPs, of the 99 KZFPs that have ChIP-exo data 89 KZFPs preferentially bind ERVs (Fig. S7B). We further selected 34 KZFPs that are significantly expressed in PriSC (Fig. S7C). However, the expression of these target ERV loci of these 34 KZFPs was not changed significantly at the mitosis-to-meiosis transition (SG to PriSC: Fig. S7D). Together with the mouse analysis, our results suggest that meiotic KZFPs are not associated with changes in ERV suppression in meiosis in mammals.

### Meiotic sex chromosome inactivation antagonizes the coevolution of KZFPs and ERVs in mammals

During meiosis, sex chromosomes undergo MSCI, which is an essential process in the male germline (Turner 2015; Alavattam et al. 2021). MSCI silences expression of most of the sex chromosome-linked genes during the pachytene and diplotene stages and largely continues to postmeiotic silencing, with a subset of genes associated with spermiogenesis escaping postmeiotic silencing (Namekawa et al. 2006; Mueller et al. 2008). Previous studies revealed that MSCI impacted the evolution of the sex chromosomes (Potrzebowski et al. 2008). For example, X chromosomes are enriched with the genes expressed before meiosis (Khil et al. 2004) and with genes that escape postmitotic silencing in spermatids (Sin et al. 2012; Mueller et al. 2013). Thus, we sought to determine how MSCI impacted the coevolution of KZFPs and ERVs in mammals. Because we found that spermatogonia-enriched KZFPs regulate meiotic ERVs, we examined the chromosome distribution of these KZFPs and ERVs. Notably, meiotic ERVs, which are targets of spermatogonia-enriched KZFPs, are enriched on autosomes and underrepresented on the sex chromosomes in mice (Fig. 7A, B), a. tendency also observed in humans (Fig. 7C, D). We next examined the chromosomal distribution of ERVs targeted by meiotic KZFPs and found that these ERVs are mostly on autosomes and underrepresented on the sex chromosomes (Fig. S8A-D). In humans, although PriSC- enriched KZFPs were not located on the X chromosome, the presence of SG-enriched KZFPs was observed on the X chromosome (Fig. S8E, F). This distribution pattern is consistent with the enrichment of genes expressed before meiosis on the X chromosome (Khil et al. 2004). These analyses suggest that MSCI antagonizes the coevolution of KZFPs and ERVs in the sex chromosomes in mammals (Fig. 7E).

**Figure 7.**
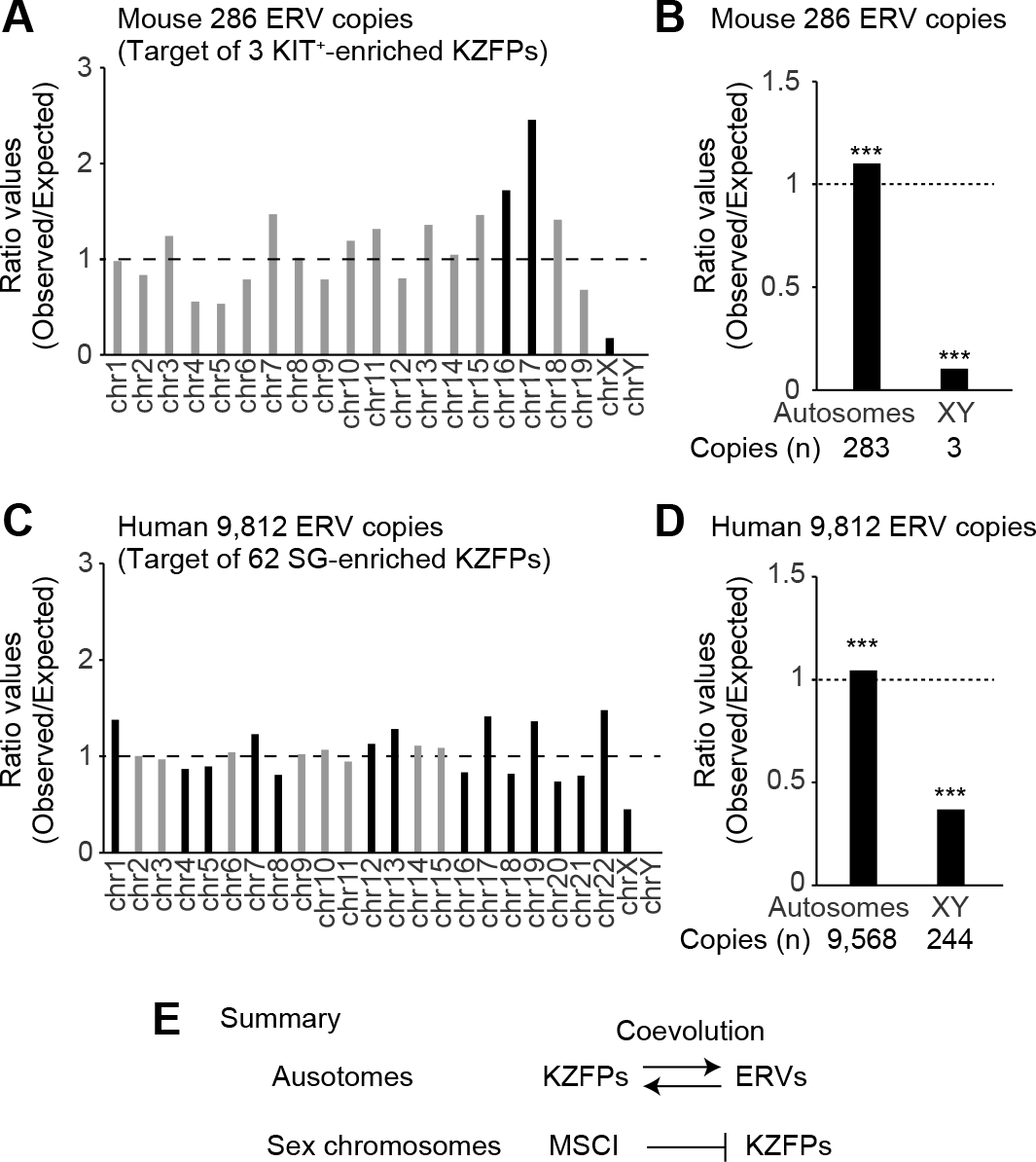
Chromosomal distribution of ERVs targeted by mouse and human spermatogonia-enriched KZFPs. **A**, Chromosomal distribution of mouse ERV copies targeted by KIT^+^-enriched KZFPs. The bar plot represents the ratio values of the observed copy number of ERVs versus the theoretically expected number. The color of each bar represents the statistical significance (Black: p<0.05, Grey: not significant, binomial test). **B**, Distribution of mouse ERV copies targeted by KIT^+^-enriched KZFPs on autosomes or sex chromosomes. Binomial test. ***: p<0.001. **C**, Chromosomal distribution of human ERV copies targeted by SG-enriched KZFPs. The bar plot is represented as same as panel A. **D**, Distribution of human ERV copies targeted by SG-enriched KZFPs on autosomes or sex chromosomes. The bar plot is represented as same as panel B. **E**, Summary. MSCI antagonizes the coevolution of KZFPs and ERVs on the sex chromosomes in mammals

## Discussion

In this study, by focusing on a family of rapidly evolving repressive transcriptional factors, KZFPs (Tadepally et al. 2008; Yang et al. 2017; Bruno et al. 2019; Wolf et al. 2020; Senft and Macfarlan 2021), we describe a molecular logic by which a large number of ERVs are controlled and specifically activated after the mitosis-to-meiosis transition. The expression of KZFPs drastically changes at the mitosis-to-meiosis transition during spermatogenesis in both in mouse and human and these changes accompany expression changes of a variety of TEs. At the ERV loci targeted by spermatogonia-enriched KZFPs, active enhancers likely formed after entering meiosis, as KZFPs were lost and H3K9me3 decreased, and H3K27ac increased. (Fig. 4H).

Retrotransposons require transcription to propagate in the germline. Thus, retrotransposons become activated in the germline where their expression is tightly controlled to minimize genome instability. What remains largely unknown is when retrotransposons are activated in the germline. Our study suggests that the time is entry into meiosis and serves as the main battlefield of rapidly expanding TEs and rapidly evolved suppressive KZFPs because their reciprocal pattern of expression of KZFPs and their target ERVs is consistent with coevolution between them. A consequence of such co-evolution would be lineage-specific germline transcriptomes. Coevolution of KZFPs and ERVs was also suggested in the context of mouse and human embryonic development (Pontis et al. 2019; Seah et al. 2019; Pontis et al. 2022). Thus, there could be several windows in the germline cycle to drive genomic evolution.

We also sought to understand the functions of KZFPs that are enriched in the meiotic stage.

Although some PS-enriched KZFPs showed a binding preference for ERVs or SINEs, we did not find any association with gene expression changes. Therefore, meiotic KZFPs may have functions other than transcriptional suppression. For example, PRDM9, a well-conserved and the most ancient type of KZFP functions as the determinant of meiotic DNA double-strand break during meiotic recombination but is not involved in gene regulation. Although human and mouse PRDM9 protein has a KRAB-A domain, it does not interact with an essential regulator of heterochromatin, KAP1, and thus is not involved in transcriptional suppression (Baudat 2010; Myers S 2010; Patel et al. 2016; Imai et al. 2017). Moreover, a previous study revealed that an evolutionally old KZFP group, termed variant KZFPs, does not interact with KAP1 and may have different roles than KAP1-dependent TE suppression (Helleboid et al. 2019).

The study identified 35 human KZFPs as variant KZFPs, and there are 26 orthologs in the mouse genome. Notably, among 26 orthologous genes, 9 KZFP genes (*Zfp202, Zkscan5, Zkscan6, Zkscan17, Zfp212, Zfp786, Zfp473, Zfp446, Zfp78*) are classified as PS-enriched KZFPs in our current study and therefore may not act as transcriptional repressors.

We find that KZFPs-targeted ERVs are underrepresented on the X chromosome, presumably due to MSCI. Intriguingly, autosomes carry many retrotransposed genes that originated from X-chromosomes to avoid MSCI; these genes are expressed later during spermatogenesis (Emerson et al. 2004). These cases of retrotranspositions coincide with the timing of dynamic changes in expression of KZFP and retrotransposons we report here. Therefore, extensive genomic evolution may, in part, be driven by MSCI at the mitosis-to-meiosis transition, where extensive epigenetic programming takes place. During meiosis, H3K9me3 is established on the X chromosomes by SETDB1, but this action of SETDB1 is regulated downstream of the DNA damage response (DDR) pathway, which is the master regulator of MSCI (Hirota et al. 2018). Therefore, both X-linked protein-coding genes and TEs are also highly enriched with H3K9me3 in pachytene spermatocytes (Liu et al. 2019), likely by the action of DDR-directed H3K9me3, which can be KZFP-independent. Of note, we previously showed that enrichment of ERV enhancers marked with H3K27ac on the X chromosome in pachytene spermatocytes (Sakashita et al. 2020). One possible reason for this observation is that these X-linked ERVs may not require KZFPs as a regulatory mechanism. Indeed, in addition to H3K9me3, H3K27ac is established by the DDR pathway on the sex chromosomes in meiosis (Adams et al. 2018). Our observation suggests that distinct mechanisms regulate the activities of ERVs on autosomes and sex chromosomes during spermatogenesis.

Our study used the binding site information generated in previous studies (Imbeault et al. 2017; Wolf et al. 2020) and focused on analyses of a limited number of KZFPs. However, we predict that not only is there a much broader network of interactions taking place between KZFPs and ERVs in the mammalian male germline but also that germline functions of KZFPs are much broader than previously thought, e.g., the cases for PRDM9 and other KZFPs required for imprinting control (Li et al. 2008; Takahashi et al. 2019). Our study provides a foundation to pursue functional studies and genomic studies for mammalian evolution.

Our results provide the first evidence that an evolutionary arms race between virus-derived sequences, ERVs, and endogenous defensive machinery, KZFPs, takes place during the mitosis-to-meiosis transition in germ cells. Expression of KZFPs mirrors expression of ERVs, and the target ERVs bound by spermatogonia-enriched KZFPs are highly covered with H3K9me3 in spermatogonia. After germ cells enter meiosis, H3K9me3 disappears and the ERV loci acquire an active enhancer mark H3K27ac mark, accompanied by the loss of KZFPs, leading to activation of adjacent genes in meiotic pachytene spermatocytes. This trend is found in mice and humans, suggesting that mammalian species developed a regulatory machinery composed of rapidly evolved KZFPs and ERV activity in the male germline. In addition, KZFPs-dependent ERV suppression mainly operates on autosomes and is underrepresented on the sex chromosomes. We propose that epigenetic programming in the mammalian germline during the mitosis-to-meiosis transition facilitates coevolution of KZFPs and TEs on autosomes and is antagonized by MSCI.

## Methods

### RNA-seq data analysis

Raw reads of previous RNA-seq reads were first trimmed using Trim Galore (https://github.com/FelixKrueger/TrimGalore) (version 3.3) and then aligned to either the mouse (GRCm38/mm10) or human (GRCh38/hg38) genomes using STAR alignment software (Dobin et al. 2013) (version 2.5.3a). Multiple mapped reads were used to quantify gene expression, whereas only uniquely aligned reads were used to quantify TE expression. All aligned reads were then sorted and indexed by SAMtools (Li et al. 2009) (version 1.11) function sort and index, respectively. The gene annotation file was obtained from the GENCODE database (v41 for humans, and vM25 for mice, https://www.gencodegenes.org). The list of KZFPs was obtained from the previous study (Imbeault et al. 2017). TEs and repeat sequence annotation files were downloaded from the UCSC genome browser RepeatMasker Track (https://genome.ucsc.edu/cgi-bin/hgTrackUi?g=rmsk), which is based on the Repbase (Bao et al. 2015) library. After quantification, unexpressed TE copies (<Raw read counts: 2) through data sets were removed. To determine differentially expressed genes and TE copies, read counts were used as input data for the DESeq2, the R package (Love et al. 2014) (version 1.34.0). To perform k-means clustering analysis for the subset of genes and TEs and draw heatmaps, we used the web tool Morpheus distributed by the Broad Institute (https://software.broadinstitute.org/morpheus). To perform

GO enrichment analysis for the subset of KZFPs, we used the web tool DAVID (Huang da et al. 2009; Sherman et al. 2022).

### ChIP-seq and ChIP-exo data analysis

Raw reads of previous ChIP-seq and ChIP-exo reads were first trimmed using Trim Galore and then aligned to either the mouse (GRCm38/mm10) or human (GRCh38/hg38) genomes using bowtie2 software (Langmead and Salzberg 2012) (version 2.4.4). All aligned reads were then sorted and indexed by SAMtools function sort and index, respectively. Peak calling for ChIP-seq and ChIP-exo data was performed using MACS2 (Zhang et al. 2008) (version 2.2.7.1) following the setting described in the previous study (Imbeault et al. 2017; Wolf et al. 2020). The expected background was estimated by randomly generating and calculating numbers of background genomic regions equal to the numbers of ChIP-seq peak regions determined by BEDTools (Quinlan and Hall 2010) (version 2.30.0) function random.

Target TEs of each KZFP were determined by examining the overlap between TE annotations and detected peaks by BEDTools function intersect. The normalized frequency of H3K9me3 and H3K27ac was calculated by BEDTools function coverage. The genomic annotation of target TEs was analyzed by HOMER software (Heinz et al. 2010) (version 4.11) function annotatePeaks.pl using a genome coordinate file (bed file) which contains the genomic location of target ERVs as input data.

The program deepTools (Ramirez et al. 2016) (version 3.5.1) was used to draw tag density plots and heatmaps for read enrichment (H3K9me3, H3K27ac and each KZFP ChIP-seq reads). To detect genes adjacent to target ERVs, we used the ChIPpeakAnno, the R package (Zhu J Lihua 2010) (version 3.28.1) using a genome coordinate file (bed file) which contains the genomic location of target ERVs as input data. To visualize read enrichment over representative genomic loci, bigwig files were created from sorted BAM files using deepTools function bamCoverage, and visualized by using IGVTools (Robinson T James 2011). To determine the evolutional age of target TEs, we referred to the database of the research community Dfam (Storer et al. 2021).

### ES cell culture and KZFP knockdown experiment

The J1 male ESCs were cultured in ESC medium (15% FBS (Hyclone, Japan), 1x GlutaMAX (Invitrogen, MA), 1x MEM non-essential amino acids solution (Invitrogen), 1x penicillin-streptomycin (Invitrogen), and 55 μM β-mercaptoethanol in Knockout D-MEM (Invitrogen) containing 2i (1 μM PD0325901, LC Laboratories, MA; and 3 μM CHIR99021, LC Laboratories) and LIF (1,300 U /ml, Merck, MA) on cell culture plates coated with 0.2% gelatin under feeder-free conditions.

One day before transfection, 1.0×10^5^ cells were seeded onto a 24-well plate coated with 0.2% gelatin. For transfection, Lipofectamine RNAiMAX (Invitrogen) was used following the manufacturer’s guidelines with 10 pmol siRNA. 6 days after transfection, cells were lysed and total RNA was isolated by using RNeasy Plus Micro Kit (Qiagen, Netherlands). 100ng of total RNA was reverse transcribed by using SuperScript IV (Invitrogen). cDNA was subjected to qRT-PCR using PowerUp SYBR Green Master Mix (Applied Biosystems, MA) with primers listed in Supplemental Table 3.

### Evaluation of sequence similarities across mammalian species

We sought to calculate sequence similarities and detect orthologous adjacent genes to mouse and human target ERVs across mammalian species. To do so, we applied a list of mouse and human ERV-adjacent genes to BioMart (Kinsella et al. 2011) to compute sequence similarities, that is, percent identities of target genes in other species in comparison to the respective mouse query genes. The divergence time of each evolutionary clade was determined by referring to the web tool Timetree (Kumar et al. 2022).

### Statistical analyses

All the statistical analyses were performed by R software (https://www.r-project.org) (version 4.1.3).

### Availability of Data

RNA-seq raw reads from previous studies were downloaded under the accession numbers as follows:

| Species | Material | GEO accession | SRA study accession |
| --- | --- | --- | --- |
| mouse | Brain | GSE93496 | SRP013027 |
| mouse | Heart | GSE93483 | SRP013027 |
| mouse | Kidney | GSE93492 | SRP013027 |
| mouse | Liver | GSE93485 | SRP013027 |
| mouse | Lung | GSE93502 | SRP013027 |
| mouse | Ovary | GSE93494 | SRP013027 |
| mouse | Placenta | GSE90203 | SRP013027 |
| mouse | Spleen | GSE93503 | SRP013027 |
| mouse | Testis | GSE93488 | SRP013027 |
| mouse | Sertoli | GSE154484 | SRP272065 |
| mouse | E18.5PSG | - | ERP015330 |
| mouse | THY1<sup>+</sup> | GSE55060 | SRP037992 |
| mouse | KIT<sup>+</sup> | GSE89502 | SRP092547 |
| mouse | PS | GSE55060 | SRP037992 |
| mouse | RS | GSE55060 | SRP037992 |
| mouse | Sperm | - | DRP000508 |
| human | SG | - | SRP075318 |
| human | PriSC | - | SRP075318 |
| human | RS | - | SRP075318 |

ChIP-seq and ChIP-exo raw reads from previous studies were downloaded under the accession numbers as follows:

| Species | Material | ChIP-target | GEO accession | SRA study accession |
| --- | --- | --- | --- | --- |
| mouse | B6 ES cells, F9 EC cells | murine KZFPs | GSE115291 | SRP149699 |
| mouse | KIT<sup>+</sup> | H3K9me3 | GSE137744 | SRP222567 |
| mouse | PS | H3K9me3 | GSE137744 | SRP222567 |
| mouse | KIT<sup>+</sup> | H3K27ac | GSE107398 | SRP125760 |
| mouse | PS | H3K27ac | GSE107398 | SRP125760 |
| mouse | TypeB spermatogonia | H3K9me3 | GSE132446 | SRP200968 |
| mouse | Leptonema | H3K9me3 | GSE132446 | SRP200968 |
| mouse | Mid-Zygonema | H3K9me3 | GSE132446 | SRP200968 |
| mouse | Mid-pachynema | H3K9me3 | GSE132446 | SRP200968 |
| mouse | Diplonema | H3K9me3 | GSE132446 | SRP200968 |
| mouse | TypeB spermatogonia | H3K27ac | GSE132446 | SRP200968 |
| mouse | Leptonema | H3K27ac | GSE132446 | SRP200968 |
| mouse | Mid-Zygonema | H3K27ac | GSE132446 | SRP200968 |
| mouse | Mid-pachynema | H3K27ac | GSE132446 | SRP200968 |
| mouse | Diplonema | H3K27ac | GSE132446 | SRP200968 |

Single-cell RNA-seq data from the previous study (Hermann et al. 2018a) was downloaded from the Mendeley Data Deposition (https://data.mendeley.com/datasets/kxd5f8vpt4/1), and was processed using 10x Genomics Loupe Browser (version 6.0.0, https://www.10xgenomics.com/products/loupe-browser).

## Author contributions

K.O. and S.H.N. designed the study. K.O., A.S., designed and interpreted the computational analyses. All authors interpreted the results. K.O, R.M.S., and S.H.N. wrote the manuscript with critical feedback from all other authors. S.H.N. supervised the project.

## Acknowledgments

We thank the members of the Namekawa laboratory for discussion and helpful comments regarding this manuscript, Todd Macfarlan for providing key unpublished data for analysis, and Neil Hunter for discussion.

