## SupplementaryFigures for "KRAB-zinc-finger proteins regulate endogenous retroviruses to sculpt germline transcriptomes and genome evolution"

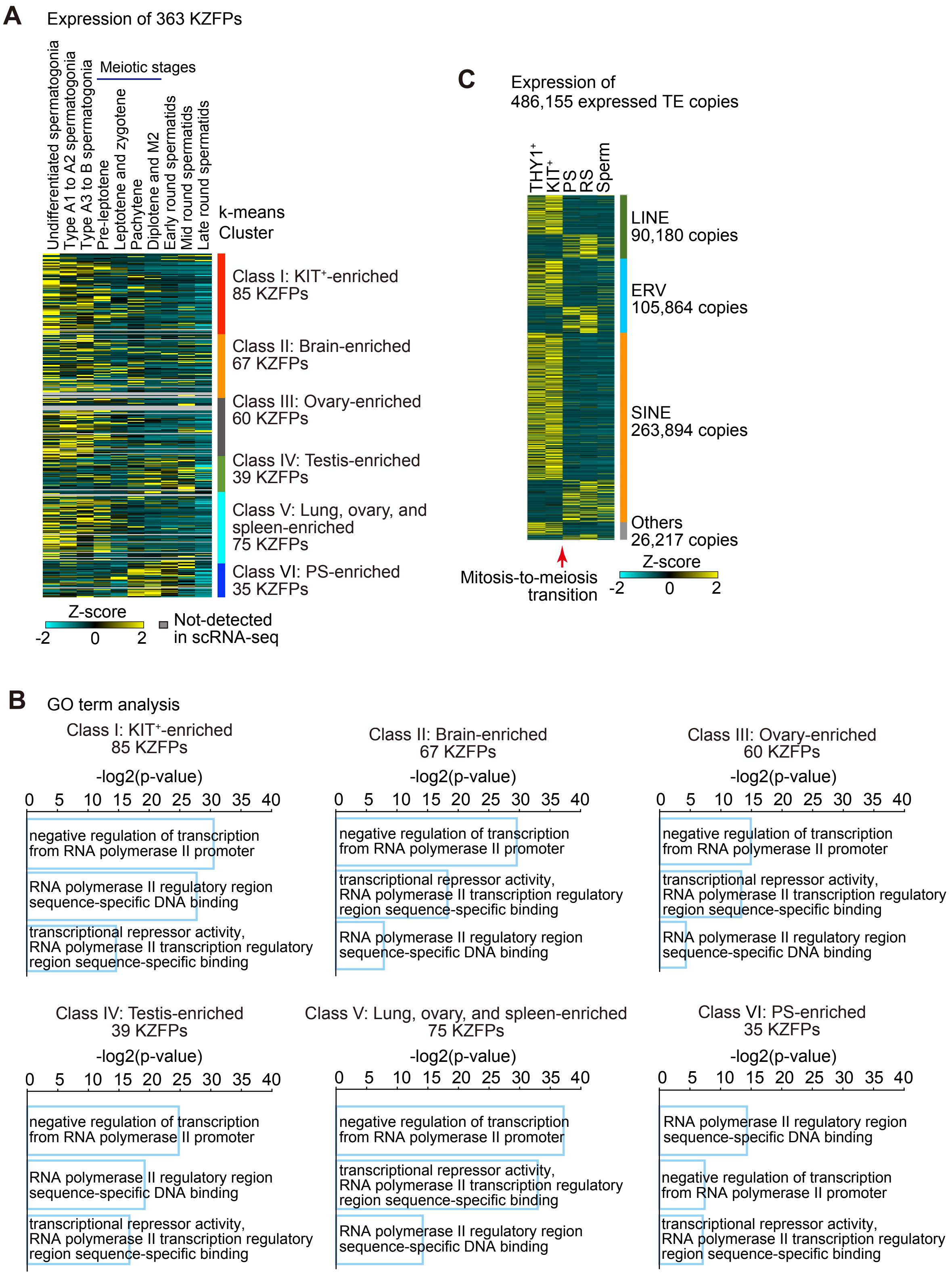


**Fig. S1: Expression of KZFPs and TEs during mouse spermatogenesis and GO term analyses.**

**A**, A heatmap showing the expression (single-cell RNA-seq) for KZFPs analyzed in Fig. 1B in various tissues and testicular cell types in mice. Data from adult testis were classified into 10 substages of spermatogenic cells. The expression level was calculated as the median-normalized average at the specific substage.

**B**, GO term of each KZFP cluster. The bar plot represents the -log2 p-value of each GO term.

**C**, A heatmap showing the expression (RNA-seq data) of all detected TE copies in male germ cells at the representative five stages.

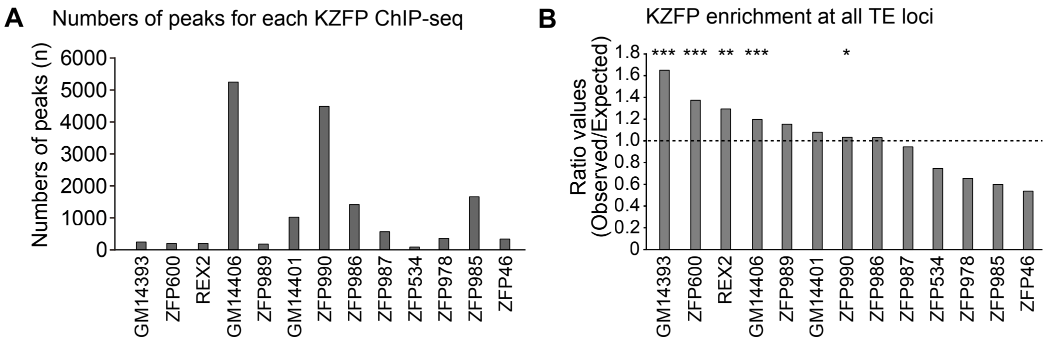


**Fig. S2: ChIP-seq analyses of KIT$^{+}$ spermatogonia-enriched KZFPs in mice**
**A**, Numbers of all detected ChIP-seq peaks.
**B**, Binding preference of KZFPs at all transposable elements (TEs). The bar plot represents the ratio values of the observed peak number versus the theoretically expected number. Binomial Test. ***: $p<0.001$, **: $p<0.01$, *: $p<0.05$.

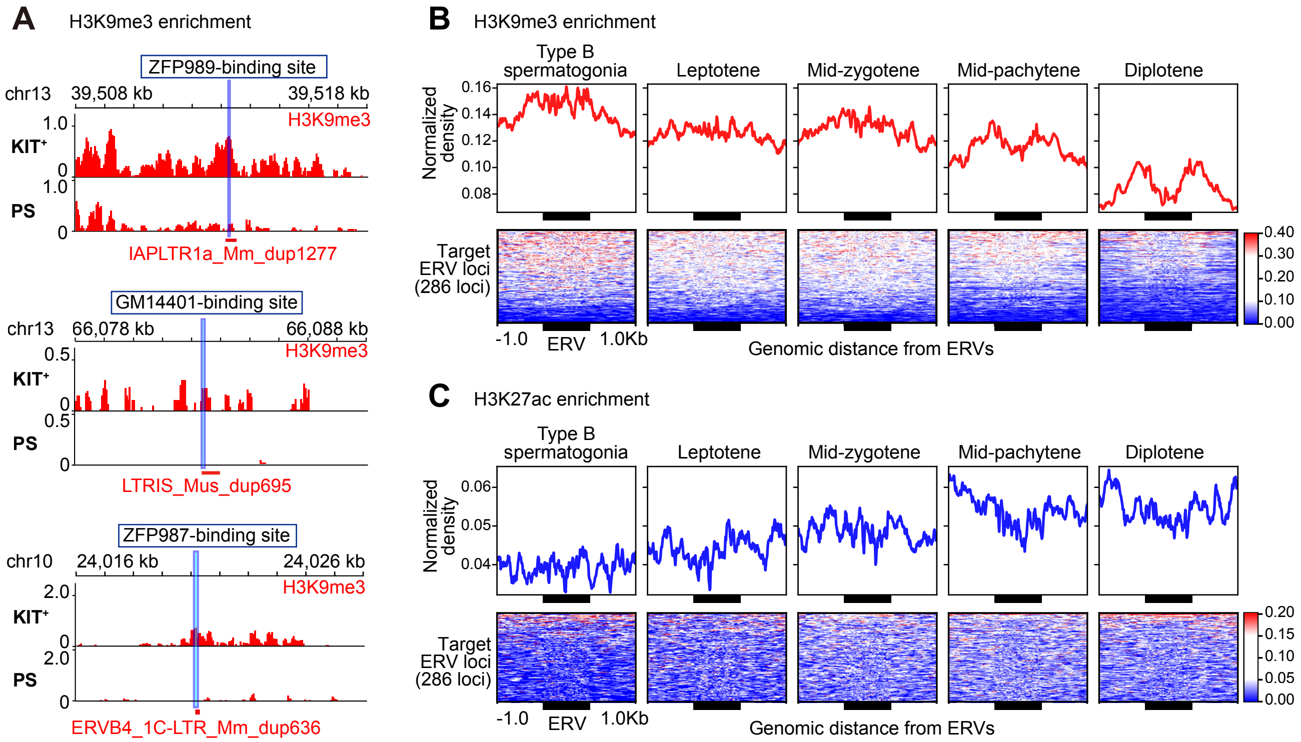


**Fig. S3: H3K9me3 and H3K27ac enrichment at the KIT⁺ spermatogonia-enriched KZFP loci in mice**

**A**, Track view of H3K9me3 enrichment at a binding site of ZFP989 (Top), GM14401 (Middle), and ZFP987 (Bottom) in KIT⁺ spermatogonia and PS.

**B, C,** The top panel represents an average density of H3K9me3 (**B**) and H3K27ac (**C**) at the target ERVs ± 1kb region. The bottom heatmap represents normalized H3K9me3 signal intensity.

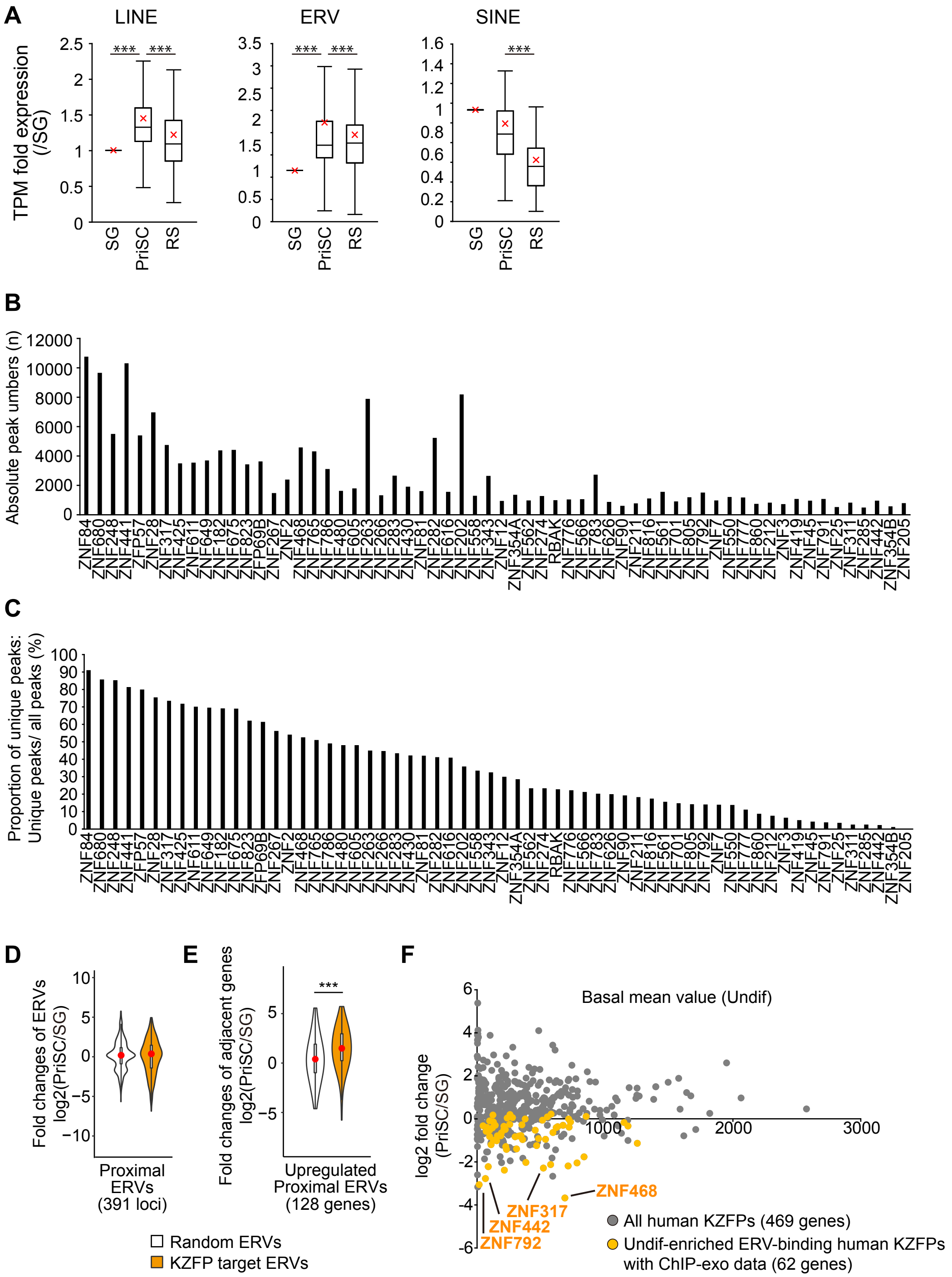


**Fig. S4: ChIP-exo analyses of SG-enriched KZFPs in humans**.

**A**, Relative expression (RNA-seq data) of each TE class during human spermatogenesis. The values of SG were set to 1. The red crosses represent the average values. ANOVA followed by Tukey-HSD test. ***: p<0.001.

**B**, The bar plot represents the numbers of peaks from previous ChIP-exo data (Imbeault et al. 2017). 62 KZFPs enriched in SG are listed.

**C**, Unique peak frequency of SG-enriched KZFPs. The bar plot represents the percentage of uniquely observed peaks.

**D**, Fold-expression change of target proximal ERVs at the mitosis-to-meiosis transition (PriSC/SG), Student' s t-test, not significant.

**E**. Fold-expression change of the genes adjacent to upregulated proximal ERVs at the mitosis-to-meiosis transition (PriSC/SG). Student' s t-test. ***: p<0.001.

**F**, Expression and fold change of human KZFPs at the mitosis-to-meiosis transition. 4 KZFPs that showed the most drastic change (ZNF792, ZNF442, ZNF317, and ZNF468) were chosen for further analyses as shown in Fig. 5G.

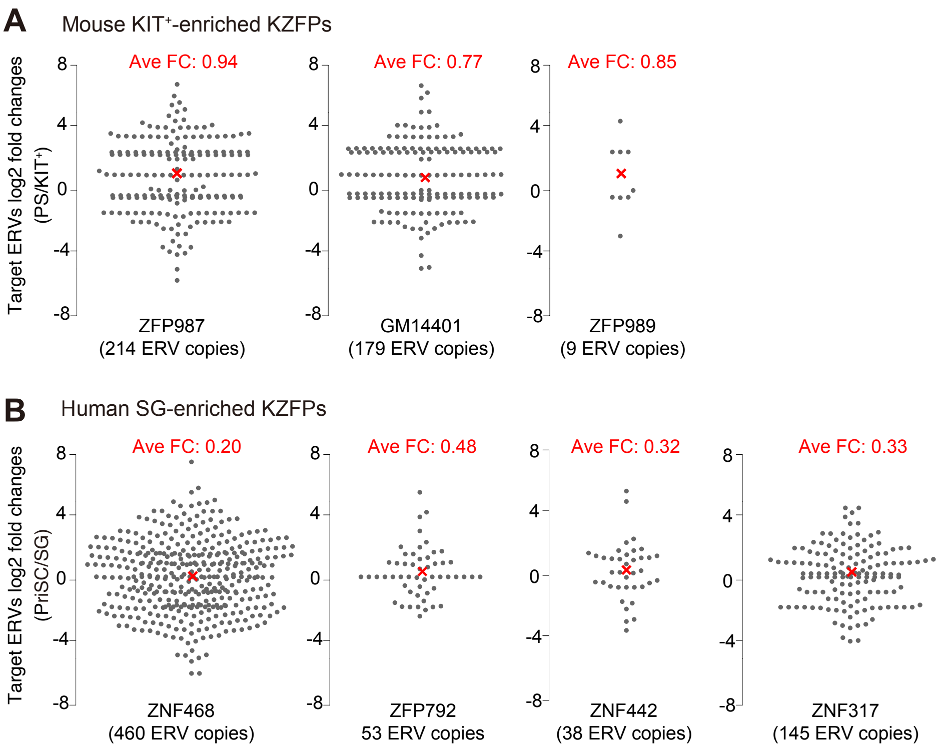


**Fig. S5: Expression change of spermatogonia-enriched KZFPs at the mitosis-to-meiosis transitions in mice and humans.**

**A, B,** Log2-fold changes of ERVs targeted by mouse (A) and human (B) spermatogonia-enriched KZFPs. The red crosses represent the average fold change values (shown on top of the panels). The numbers in parentheses represent the detected ERV copy numbers.

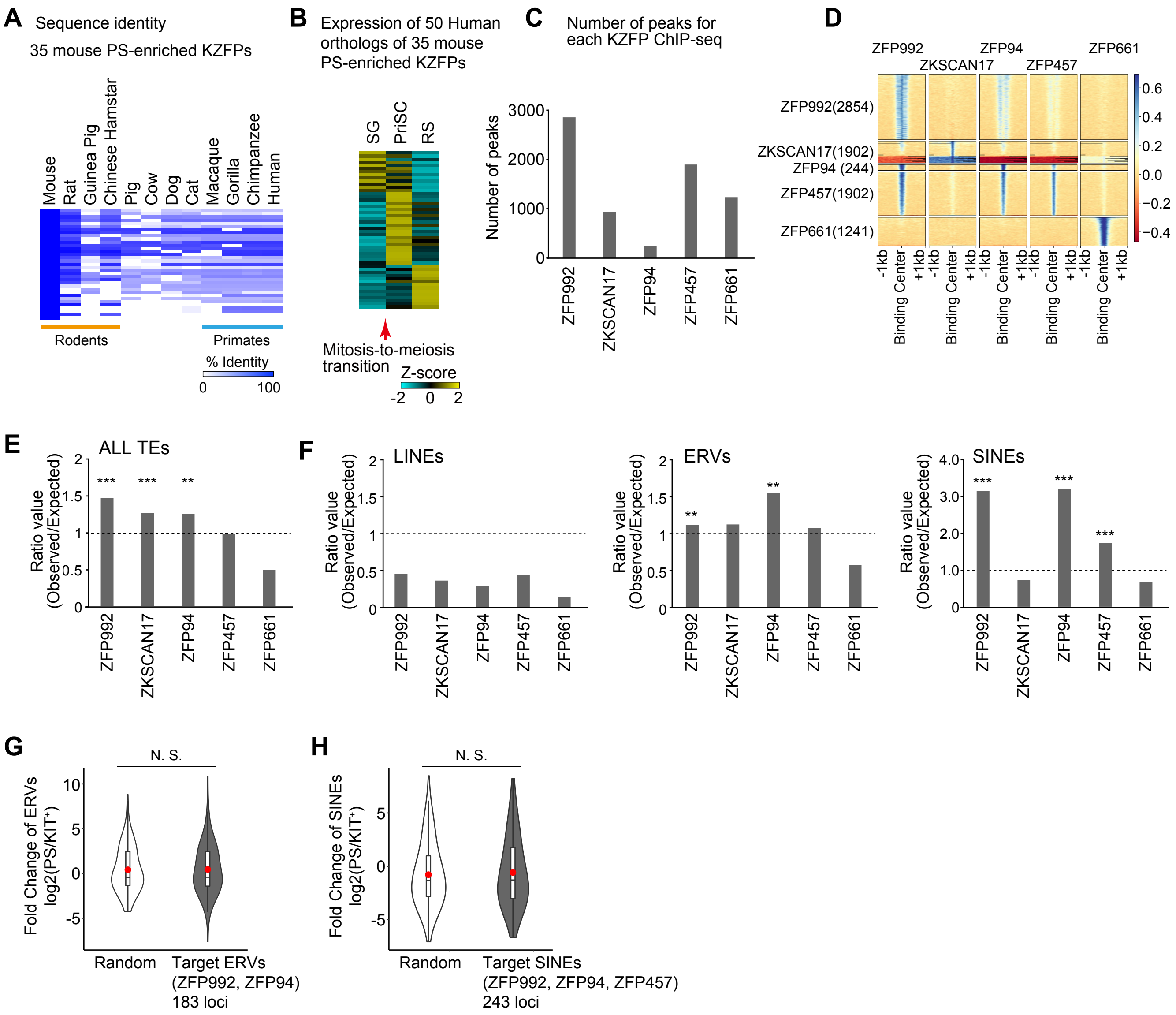


**Fig. S6: Analyses of mouse PS-enriched KZFPs**

**A**, Heatmap represents the identity of 35 murine PS-enriched KZFPs compared among mammalian species.

**B**, Heatmap represents the expression of 50 human orthologous genes.

**C**, Numbers of all detected ChIP-seq peaks.

**D**, Heatmap of the binding profiles of PS-enriched KZFPs. The numbers in parentheses represent the numbers of detected peaks.

**E, F,** Binding preference of KZFPs at all TEs (**E**) and each TE subclass (**F**). The bar plot represents the ratio values of the observed peak number versus the theoretically expected number. Binomial test. ***: $p<0.001$, **: $p<0.01$.

**G, H,** Fold-expression change of ERVs targeted by ZFP992 and ZFP94 (**G**) and SINEs targeted by ZFP992, ZFP94, and ZFP457 (**H**) at the mitosis-to-meiosis transition ($PS/KIT^{+}$), Student' s t-test. N.S.: not significant.

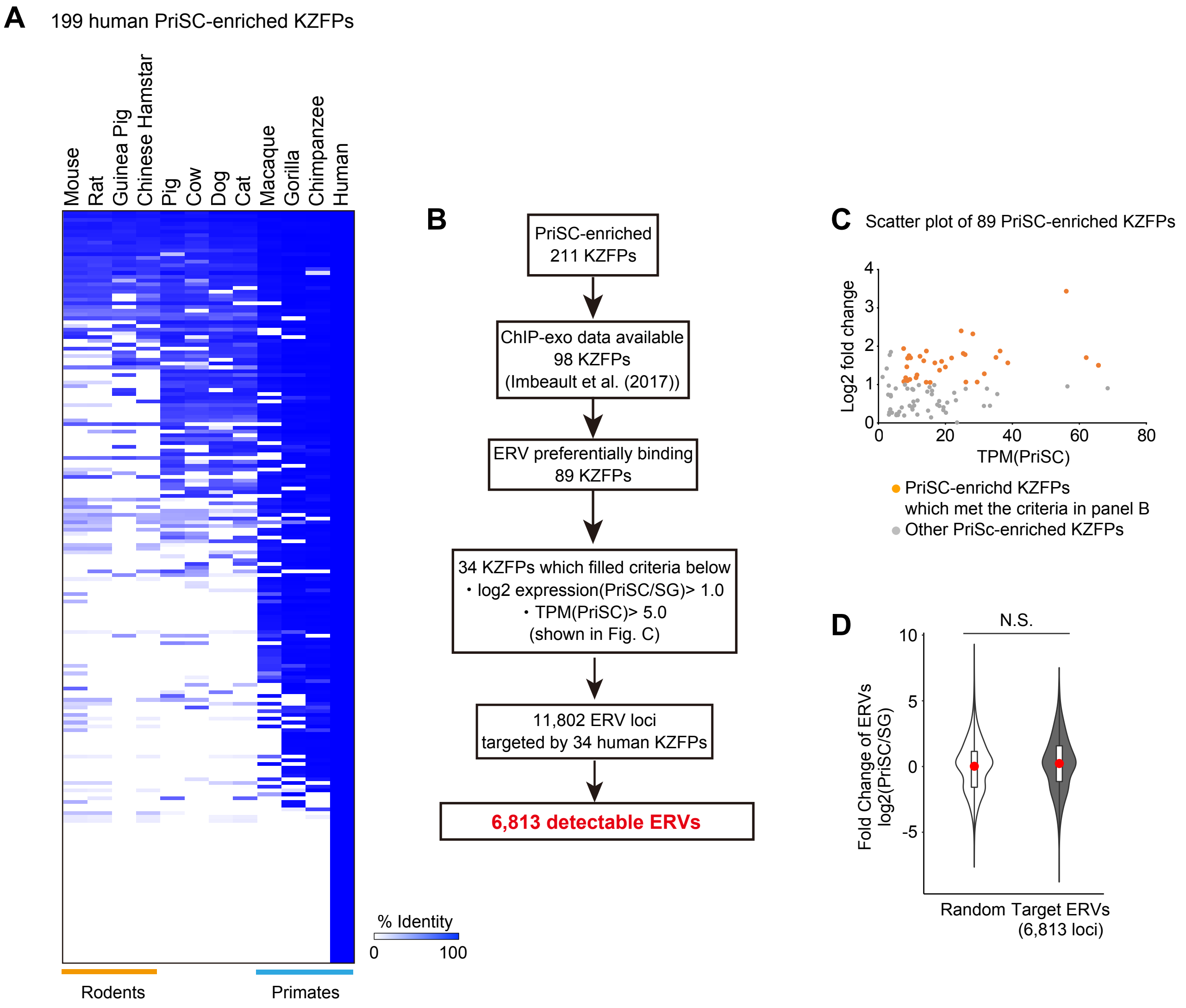


**Fig. S7: Analyses of human PriSC-enriched KZFPs**

**A**, The heatmap represents the identity of 199 human KZFPs compared among mammalian species.

**B**, Flowchart of analyses to identify target ERVs.

**C**, Scatter plot represents the TPM value (x-axis) and the log2 fold change (y-axis) of 89 PriSC-enriched KZFPs whose ChIP-exo data are available.

**D**, Fold-changes in expression of target ERVs at the mitosis-to-meiosis transition (PriSc/SG), Student’ s t-test. N.S.: not significant.

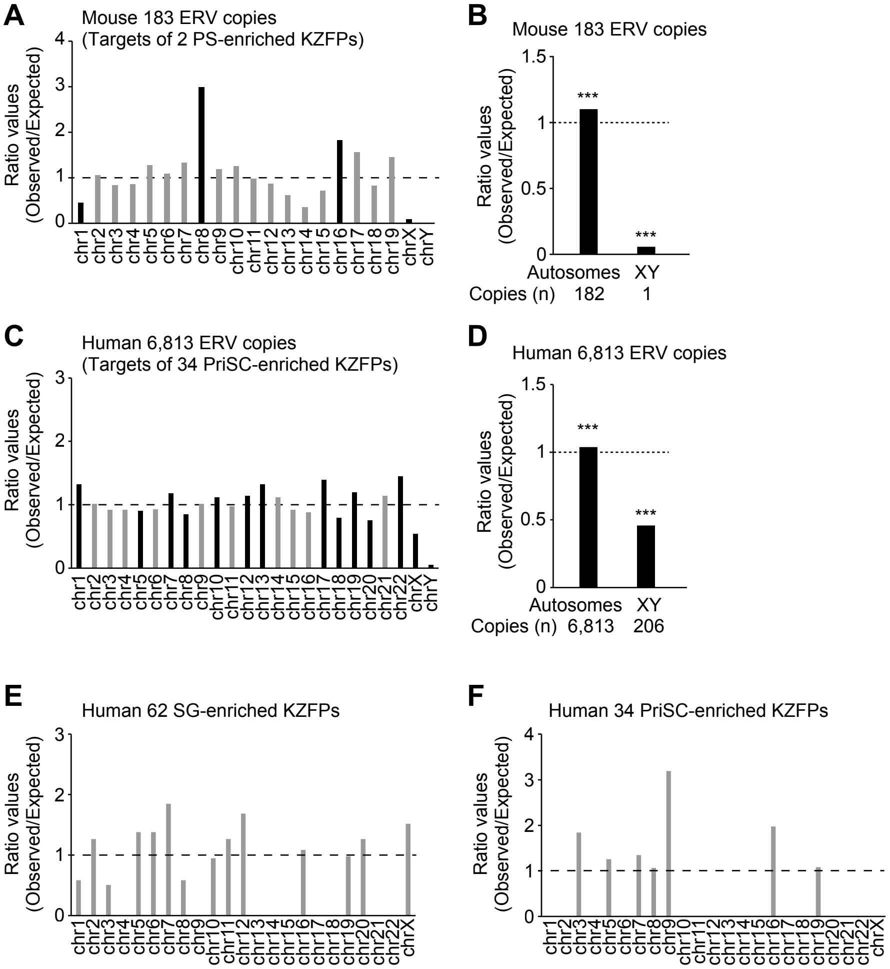


**Fig. S8: Chromosomal distributions of KZFPs and ERVs.**

**A**, Chromosomal distribution of mouse ERV copies targeted by PS-enriched KZFPs. The bar plot represents the ratio values of the observed copy number of ERVs versus the theoretically expected number. The color of each bar represents the statistical significance (Black: $p<0.05$, Grey: not significant, binomial test).

**B**, Distribution of mouse ERV copies targeted by PS-enriched KZFPs on autosomes or sex chromosomes. Binomial Test. ***: $p<0.001$.

**C**, Chromosome distribution of human ERV copies targeted by PriSC-enriched KZFPs. The bar plot is represented as same as panel A.

**D**, Distribution of human ERV copies targeted by PriSC-enriched KZFPs on autosomes or sex chromosomes. The bar plot is represented as same as panel B.

**E**, Chromosome distribution of human 62 SG-enriched KZFPs. The bar plot is represented as same as panel A.

**F**, Chromosome distribution of human 34 PriSC-enriched KZFPs. The bar plot is represented as same as panel A.
